# RosetteArray Platform for Quantitative High-Throughput Screening of Human Neurodevelopmental Risk

**DOI:** 10.1101/2024.04.01.587605

**Authors:** Brady F. Lundin, Gavin T. Knight, Nikolai J. Fedorchak, Kevin Krucki, Frank Siepel, Nisha Iyer, Jack E. Maher, Nicholas R. Izban, Abilene Roberts, Madeline R. Cicero, Joshua F. Robinson, Bermans J. Iskandar, Rebecca Willett, Randolph S. Ashton

## Abstract

Neural organoids have revolutionized how human neurodevelopmental disorders (NDDs) are studied. Yet, their utility for screening chemical hazards and prospective therapeutics for NDDs is limited by a lack of morphological reproducibility and cost-effective scalability. Here, we describe the RosetteArray platform, which can be used as an off-the-shelf, 96-well plate assay that standardizes incipient forebrain and spinal cord organoid morphogenesis as adherent, micropatterned, 3-D, singularly polarized neural rosette tissues (∼200 and ∼800 per well, respectively). Seeded directly from cryopreserved human pluripotent stem cells, RosetteArrays are cultured over 6-8 days and fixed, immunostained and imaged *in situ* to enable artificial intelligence-based quantitative analysis. By screening the inception of ∼75,000 neural organoids throughout this manuscript, we provide proof-of-concept demonstrations of the platform’s utility for detecting developmental neurotoxicity hazard and screening genetic and environmental factors known to cause clinical Neural Tube Defect risk. Given the documented perturbation of rosette morphogenesis in neural organoid models of several NDDs, the RosetteArray platform could enable quantitative high-throughput screening (qHTS) of human neurodevelopmental risk across regulatory and precision medicine applications.

## INTRODUCTION

Human pluripotent stem cell (hPSC)-derived neural organoids are prominent *in vitro* models of human central nervous system (CNS) development. Their biological relevance complements, or in some respects exceed, traditional animal models^1–3^. Formed through spontaneous self-organization of differentiating hPSC aggregates, neural organoids can recapitulate extensive aspects of CNS tissue morphogenesis and obtain organotypic levels of neural cell phenotype diversity^4–8^. As such, they are promising tools for toxicological screening, disease modeling, and drug discovery^9^. However, in the absence of bioengineered controls, their emergent 3-D morphology, cytoarchitecture, and composition can be variable, limiting inter- and intra-batch comparisons as well as scalability of post-hoc analyses^10,11^. This impedes cost-effective and scalable implementation of neural organoid technology as a quantitative high-throughput screening (qHTS) tool for detecting developmental neurotoxicity (DNT) hazards associated with chemical/drug exposures, i.e., environmental factors, and neurodevelopmental risk associated with genetic variants/mutations, i.e., genetic factors. Consequently, the multifactorial etiologies, i.e., combination of genetic and environmental factors, of various human neurodevelopmental disorders (NDDs), e.g., Neural Tube Defects (NTDs)^12,13^, Autism Spectrum Disorder^14,15^, and Schizophrenia^16^, remains unclear. Overall, this hinders progress towards effective prevention of DNT hazards and discovery of precision medicine prophylactic and therapeutic approaches for NDDs^17,18^.

Neural rosettes comprised of polarized neuroepithelial cells (NECs), a.k.a. apical radial glial cells, model the inception of CNS morphogenesis and are the morphogenetic centers of neural organoids^19,20^. Each single rosette structure is mimetic of an embryonic neural tube slice, and depending on its rostrocaudal (R/C) and dorsoventral (D/V) patterning, constituent NECs can generate the comprehensive spectrum of region-specific brain or spinal neural phenotypes^21–23^. In standard neural organoid derivation protocols, rosettes emerge from non-planar, spheroidal cell aggregates in an uncontrolled manner, producing organoids with variable rosette number, size, and shape. This strays from *in vivo* development where the neural tube emerges from a 2-D neuroectodermal plate and NECs polarize to form a single lumen. This disparity in standard protocols initiates a breakdown in reproducibility of organoid structure^23,24^.

Recently, we and others have discovered that rosette emergence at neural organoid inception can be controlled using biophysical constraints on adherent cell aggregate morphology^21,25–27^. Specifically, we integrated micropatterned culture substrates with NEC derivation protocols^28,29^ to enforce a prescribed 2-D cell monolayer morphology, which reproducibly instructs single rosette emergence upon morphing into an adherent 3-D tissue^21^. Thus, this enables one to derive microarrays of nascent neural organoids with discrete R/C regionalization, reproducible singularly polarized lumen/rosette structure, and in an adherent culture format that facilitates interrogation via microscopy, i.e., a RosetteArray. Importantly, micropatterned culture substrates can be scaled to well plate formats to theoretically enable high-throughput derivation of reproducible rosette emergence, i.e., incipient neural organoids.

As an analogue to neural tube emergence, defects in neural rosette formation or perturbed behavior of constituent NECs can be predictive of factors that cause congenital NTDs^24,30,31^, childhood-onset Autism Spectrum Disorders^14,20,32^,Schizophrenia^16^, and even adult-onset Huntington’s disease^26^ and frontotemporal dementia (FTD)/amyotrophic lateral sclerosis (ALS)^33^. Thus, since neural rosette formation requires ‘normal’ hPSC viability, proliferation, neural differentiation, and NEC physiology, *in vitro* neural rosette formation could be a predictive tool for broadly assessing DNT hazard and NDD risk factors^34^. Regarding NTDs specifically, several groups have published the use of *in vitro* neural rosette formation assays to screen for environmental and genetic risk factors. However, these assays have significant limitations. First, rosette emergence was not standardized in space and time decreasing the assays’ sensitivity^30,31,35^. Second, rosette emergence was only assessed in forebrain NECs or neural organoid cultures, whereas NTDs can occur at any point along the neural tubes’ R/C axis^24,25,30,31^. In fact, NTDs of the lower spinal cord, i.e., Myelomeningocele/ Spina Bifida, are the predominant human clinical scenario^36^. Third, none of the prior neural rosette assays were conducted using direct seeding of cryopreserved cells, which is requisite for scalable ‘off-the-shelf’ qHTS.

Here, we describe a hPSC-derived RosetteArray platform for scalable, off-the-shelf, qHTS of environmental and genetic factors, including clinically relevant multifactorial scenarios, known to cause neurodevelopmental risk. First, reproducible derivation of forebrain (FB) and spinal cord (SC) RosetteArrays from direct seeding of cryopreserved hPSCs or neuromesodermal progenitors^22,28^ (NMPs), respectively, was demonstrated. Second, the RosetteArray’s capability to detect risk associated with pharmaceuticals and agrochemicals known to cause NTDs was shown. These studies revealed differential responses between forebrain and spinal RosetteArrays to a well-known teratogen, supporting the importance of including both region-specific assays for more comprehensive coverage of DNT hazard. Third, the RosetteArray platform was evaluated for general DNT assessment in a proof-of-concept screen of a small chemical library. This included demonstrating the ability to integrate human metabolism into the assay via compound pre-digestion with S9-pooled liver fraction. Fourth, scale-up of the RosetteArray to a 96-well plate format is demonstrated with maintenance of reproducible and efficient single rosette emergence. Fifth, since disruption of folate metabolism is the best known NTD-risk factor^37,38^, RosetteArray detection of folate metabolic pathway antagonism, e.g., by Methotrexate and Dolutegravir, along with rescue by active folate supplementation was demonstrated. Lastly, gene-edited lines with mutations that perturb NTD risk-associated folate metabolic and planar cell polarity pathways^12,37^ were created. Then, the RosetteArray platform’s ability to detect both genetic and multifactorial causes for NTD risk in pathway specific and forebrain vs. spinal region-specific manners was observed. Collectively, these results support the RosetteArray platform’s potential for enabling qHTS of chemical DNT hazard and for factors that cause increased NTD risk. More broadly, they indicate that RosetteArrays could enable efficient investigation of multifactorial NDD etiology as well as screening to discover precision medicine prophylactic and therapeutic approaches.

## RESULTS

### Validation of an ‘off-the-shelf” RosetteArray assay

Previously, we demonstrated that seeding WA09 hESCs^29^ or their cervical spinal NMP progeny^22,28^ from culture onto micropatterned substrates could be used to generate arrays of singularly polarized FB and cervical SC rosette tissues (a.k.a., RosetteArrays)^21^. However, for scalable applications, an off-the-shelf protocol that uses direct seeding of cryopreserved cells is required to avoid errors caused by genetic drift during long-term cell culture^39^ and/or differences in the users’ hPSC culturing or NMP derivation techniques. Thus, we cryopreserved banks of WA09 hESCs and cervical spinal NMPs, and tested whether their direct seeding onto micropatterned substrates could also generate FB and SC RosetteArrays in a 12-well format (**Figure 1A,B**). After 24-hrs post seeding in E8 media with 10 μM Rock inhibitor (R), prospective FB and SC RosetteArrays were cultured for an additional 5 days in E6 media without^29^ or with retinoic acid (RA)^22,28^, respectively, to permit differentiation into PAX6^+^/N-cadherin^+^ (CDH2) NECs with subsequent rosette emergence (**Figure 1C**). As previously optimized^21^, FB and SC RosetteArrays were derived using micropatterned culture substrates presenting an array of 250 μm and 150 μm diameter circles, respectively, to preferentially induce single rosette emergence with regionally distinct tissue morphologies (**Figure 1D**). Post fixation and immunostaining, confocal Z-stacks of arrayed rosette tissues were collected and assessed for the number of DAPI^+^ cells and the percentage of PAX6^+^ cells (NECs) in the stack’s middle slice using CellProfiler as previously described^21^. The percentage of tissues displaying a single polarized rosette structure was quantified manually. To avoid needing to image/analyze every micropatterned tissue per well, retrospective analysis of FB RosetteArray data was used to determine that quantification of ∼40-50 tissues per well is sufficient to estimate accurate and precise results for the entire well (**Figure S1A-C**). Using this threshold, it was determined that FB and SC RosetteArrays derived using direct seeding of cryopreserved cells yielded single neural rosette efficiencies of 68.35 ± 15.35% and 63.89 ± 14.62%, respectively (**Figure 1E, 0.0% DMSO values**). Both values are lower than the 80-85% and 73.5% efficiencies observed previously when seeding direct from culture, respectively^21^.

**Figure 1.**
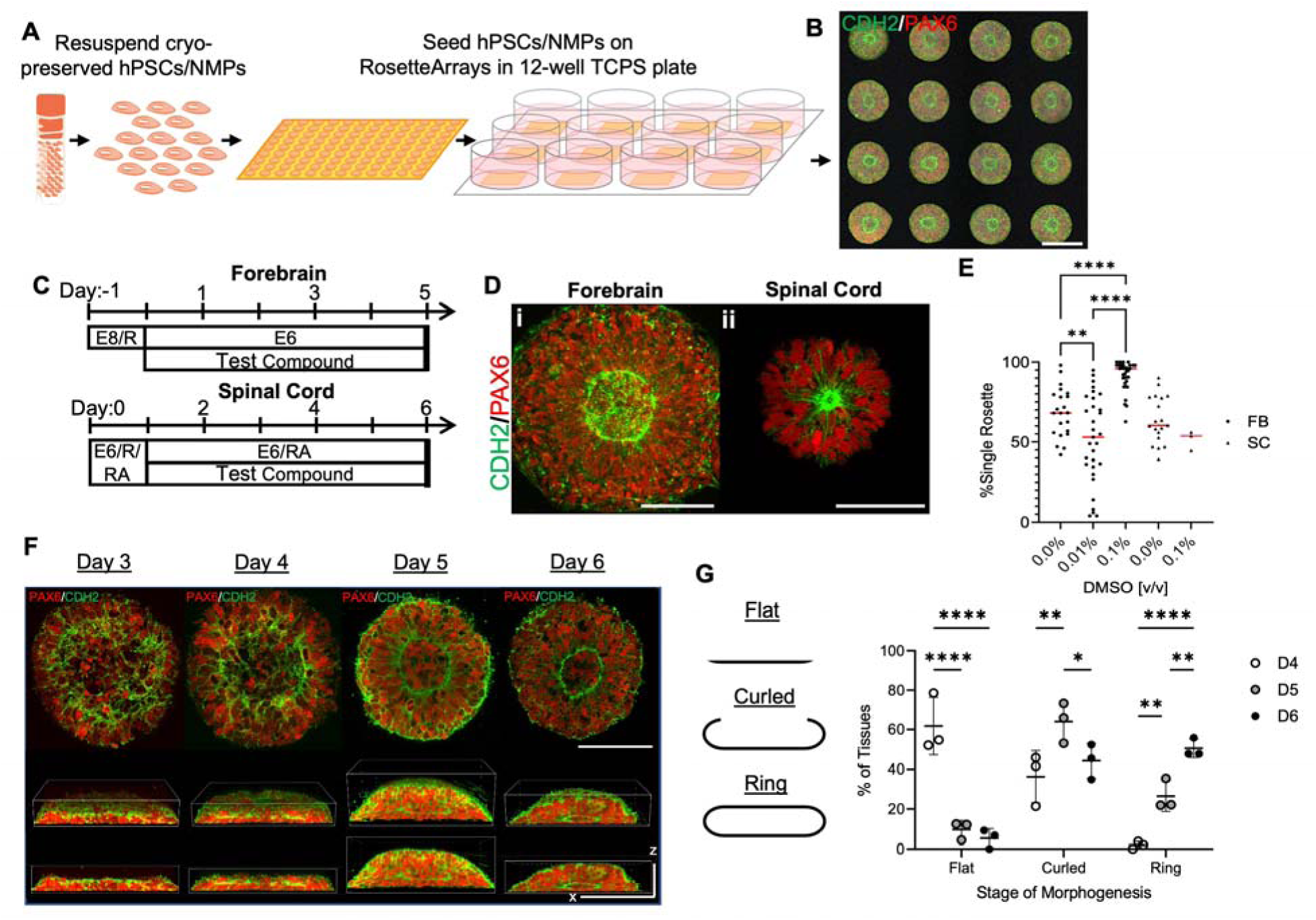
FB and SC RosetteArrays derived from cryopreserved cell banks. (A) Schematic of 12-well plate format used for direct seeding of cryopreserved WA09 hPSCs and cervical spinal NMPs onto micropatterned substrates. (B) Image of resultant immunostained FB RosetteArray showing singularly polarized (CDH2/N-cadherin^+^) neural rosette formation. (C) Culture schema for FB and SC RosetteArrays derivation with (D) images showing distinct FB (i) and SC (ii) rosette tissue morphologies. (E) Dot plot of RosetteArray derivation in E6 media supplemented with up to 0.1% v/v DMSO. Each data point is the average of a biological replicate (n=50 tissues/technical replicates per well). (F) Time course images of micropatterned rosette tissue morphogenesis at Day (D) 3-6 of FB RosetteArray derivation. Scale bars: x = 125 μm; z = 45 μm. (G) Plot tracking polarized N-cadherin ring’s profile (xz-plane) shape, i.e.,’Flat’ vs. ‘Curled’ vs. ‘Ring’, to document 2- to 3-D FB rosette tissue morphogenesis. Each data point is the average of a biological replicate, n = 40 technical replicates per well. Scale bars are (B) 250 and (D)100 µm. 1-way ANOVA with Tukey-Kramer post-hoc analysis; *p ≤ 0.05, **p ≤ 0.01, ***p ≤ 0.001, ****p ≤ 0.0001.

While concerned about the decrease in single rosette efficiency, we continued exploring the platform’s utility for chemical screening by assessing its performance in media supplemented with dimethyl sulfoxide (DMSO), a common chemical library solvent. DMSO dose response experiments using FB and cervical SC RosetteArrays were conducted to investigate the solvent’s effect on cell viability/proliferation (Cells/Tissue), neural induction^40^ (%PAX6^+^), and single rosette emergence (%Single Rosettes) (**Figure S1D-K**). Both FB and SC RosetteArrays only showed cytotoxicity and inhibition of neural induction effects at DMSO levels exceeding 1.0% v/v. However, rosette emergence was inhibited at >0.1% DMSO v/v, representing its upper limit for use in RosetteArray assays. Interestingly, 0.1% DMSO media supplementation corresponded with a significant increase in FB RosetteArray single rosette emergence (91.69 ± 8.94%) but not in SC RosetteArrays (51.45 ± 6.00%) (**Figure 1E, S3B**). We attribute this region-specific effect to DMSO’s known capacity for reversibly arresting hPSCs, but not NMPs, in the early G1 cell cycle phase thereby facilitating differentiation^41,42^. The SC RosetteArray’s persistently lower single rosette emergence efficiency remained a concern. However, collectively these results support the feasibility of deriving RosetteArrays by direct seeding of cryopreserved hPSCs and NMPs and the use of DMSO media supplementation in chemical screening, which can enhance single rosette emergence. Moreover, as demonstrated in other publications describing micropatterned morphogenesis of NEC-containing tissues^24–26^, the hPSCs are seeded as a monolayer, and coincident with neural induction (PAX6^+^/CHD2^+^), morph into 3-D hemispherical forebrain tissues with a N-cadherin^+^ polarized rosette structure by Day 5/6 of culture (**Figure 1F,G**). Further, under extended culture within a Matrigel hydrogel overlay, the arrayed FB rosette tissues demonstrate the capability to generate cortical organoids (**Supplemental Methods** & **Figure S2**).

### FB RosetteArrays detect risk of NTD-associated teratogens

NEC rosette emergence is an *in vitro* morphogenic analogue to *in vivo* neurulation. As such, we investigated whether FB RosetteArrays could detect the toxicity of compounds clinically or epidemiologically associated with NTD risk (**Figure 2**). Glycolic acid (GA) was used as a negative control^43^, and benomyl (BNM), valproic acid (VPA), methotrexate (MTX), and novobiocin (NVB) were used as positive controls. BNM has been epidemiologically correlated with NTDs and reported to antagonize the planar cell polarity (PCP) pathway, which is critical for neurulation^44,45^. VPA is an antiepileptic drug and well-known teratogen associated with NTD risk^46,47^. MTX is a chemotherapeutic and potent antagonist of folate metabolism, which is the best known NTD risk pathway^48,49^. Lastly, NVB is an antibiotic and potential chemotherapeutic that decreases the presence of extracellular matrix (ECM) protein fibronectin through inhibition of HSP-90^50^. Fibronectin plays a critical ECM role in early development and neurulation, and its NVB-mediated reduction at the neural/surface ectoderm interface was recently demonstrated to inhibit *in vitro* neural tube morphogenesis^25^.

**Figure 2.**
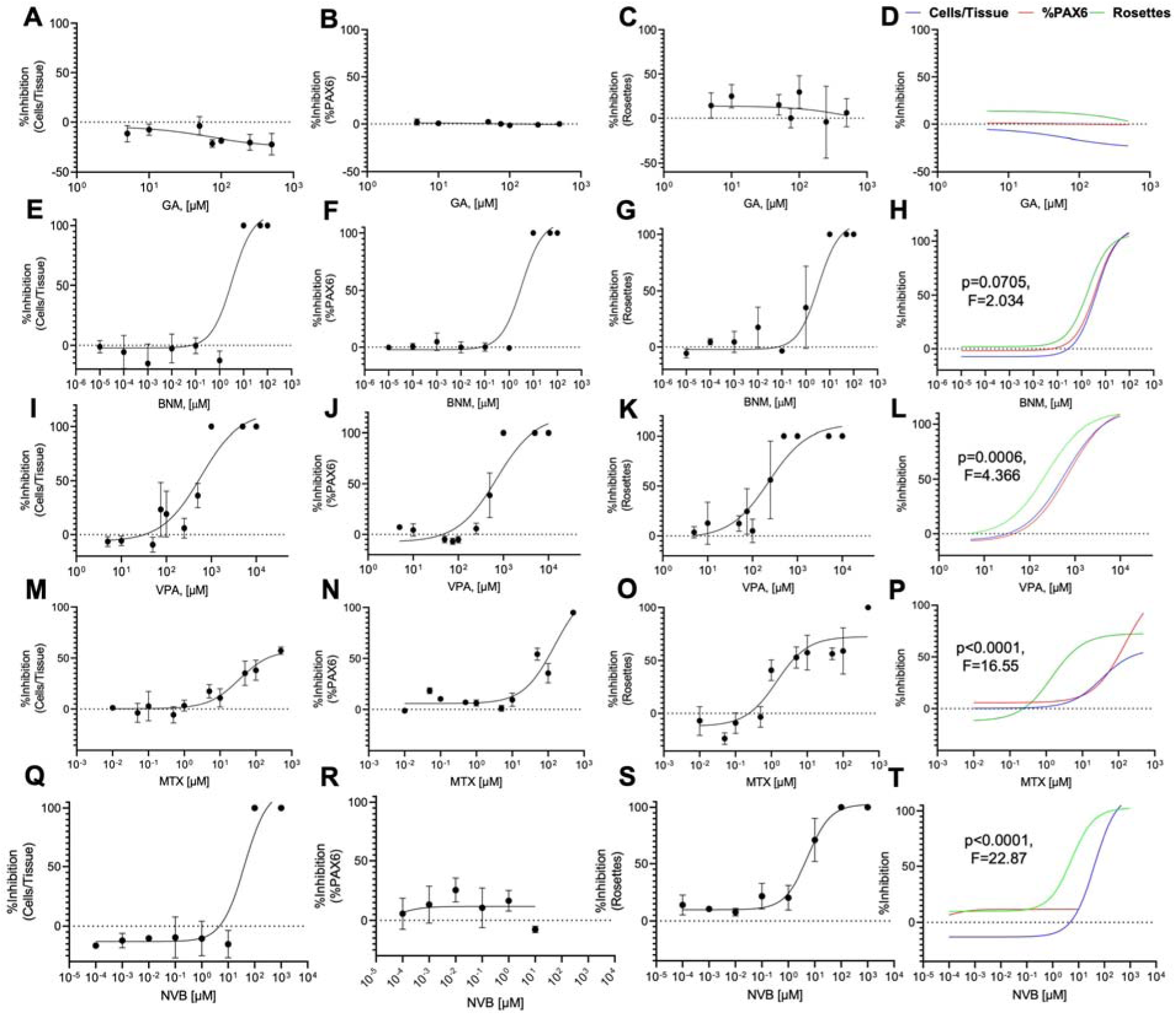
Dose-response of FB RosetteArrays exposed to Glycolic Acid, Benomyl, Valproic Acid, Methotrexate, and Novobiocin. (A) Glycolic acid dose-response graphs display percent %inhibition of (A) Cells/Tissue, (B) %PAX6 expression, and (C) single rosette emergence (Rosettes) plus (D) a composite dose-response curve overlay (Blue-Cells/Tissue, Red-%PAX6, and Green-Rosettes). This is repeated for Benomyl (E-H), Valproic Acid (I-L), Methotrexate (M-P), and Novobiocin (Q-T). Each data point is the average of a WA09 hPSC differentiation conducted in biological triplicate, n=50 technical replicates per well or 150 tissues total. Three-parameter non-linear regression used to model and compare each metric. p- and F-values indicated whether the RosetteArray metrics’ dose-response curves are better fitted with separate vs. a single curve.

NTD-risk associated chemical exposures were initiated on the first day of FB RosetteArray neural induction (Day 0) and maintained for the assay’s remaining 5 days in E6 media with or without DMSO supplementation (**Figure 1C, S3**). Juxtaposition of dose-response curves for the RosetteArray’s three primary metrics showed distinct profiles for each NTD risk compound. As expected^43^, GA did not inhibit cell any RosetteArray metrics at the levels tested (**Figure 2A-D, S3A**). However, BNM exhibited cytotoxic (Cells/Tissue) effects along with inhibition of neural induction (%PAX6) and rosette emergence (Rosettes) at >1 µM (**Figures 2E-H, S3C,G)**. Similarly, VPA and MTX exhibited cytotoxicity at the highest tested concentrations (**Figure 2I, M**), but at non-cytotoxic concentrations, they also displayed inhibition of single rosette emergence (**Figures 2I-L, M-P and S3D,E,H,I**). For example, MTX’s rosette emergence IC_50_ was ∼108-fold and ∼25-fold less than those noted for cytotoxicity and inhibition of PAX6 neural induction, respectively (**Figures 2P, S3I**). MTX’s accentuated effect on rosette emergence is indicative of its targeted inhibition of the NTD-associated folate metabolic pathway^51^. Similarly, novobiocin also inhibited rosette emergence at concentrations showing no effects on cell viability/proliferation or neural induction metrics (**Figures 2Q-T and S3F,J)**. Interestingly, the FB RosetteArray detected novobiocin’s NTD risk despite not having non-neural ectodermal cells, i.e., the purported source of fibronectin matrix production in other *in vitro* neural tube models^25^. Overall, the sensitivity of the neural rosette morphogenic metric highlights the RosetteArray’s ability to detect perturbations to pathways known to orchestrate *in vivo* neurulation and whose disruption causes NTD risk.

### RosetteArray shows regional differences in VPA’s NTD risk

FB RosetteArray assays can detect VPA’s NTD risk showing complete inhibition of rosette emergence at ≥0.5 mM concentrations (**Figure 2I-L**). However clinically, VPA exposures are predominantly observed to cause spinal NTDs^52,53^. Thus, VPA’s effect on cervical SC RosetteArrays was evaluated for comparison (**Figure 3A,B**). In contrast to FB RosetteArrays, VPA did not significantly inhibit cervical SC RosetteArray cell viability/proliferation, neural induction, or rosette emergence up to 0.5 mM (**Figure 3C-E**). Yet, the N-cadherin^+^ polarized ring area, i.e., ‘Rosette Area’, in cervical SC rosettes increased in a dose-responsive manner, which was a phenomenon not observed in VPA-exposed FB rosettes (**Figure 3B,F**). This indicates a region-specific response to VPA exposure captured by the RosetteArray platform, aligning with clinical data^52,53^ and mouse embryo studies, in which VPA exposure biomechanically disrupts closure of the posterior neural tube^47^. Concordantly, we previously showed that biomechanical differences between FB and SC rosette tissues were responsible for needing different micropattern dimensions (i.e., 250 vs. 150 μm, respectively) to induce optimal singular rosette emergence^21^. Thus, the RosetteArray platform’s ability to interrogate hPSC-derived neural rosette morphogenesis across the neuraxis may enable discrimination between adverse effects in rostral versus caudal NTD mechanisms.

**Figure 3.**
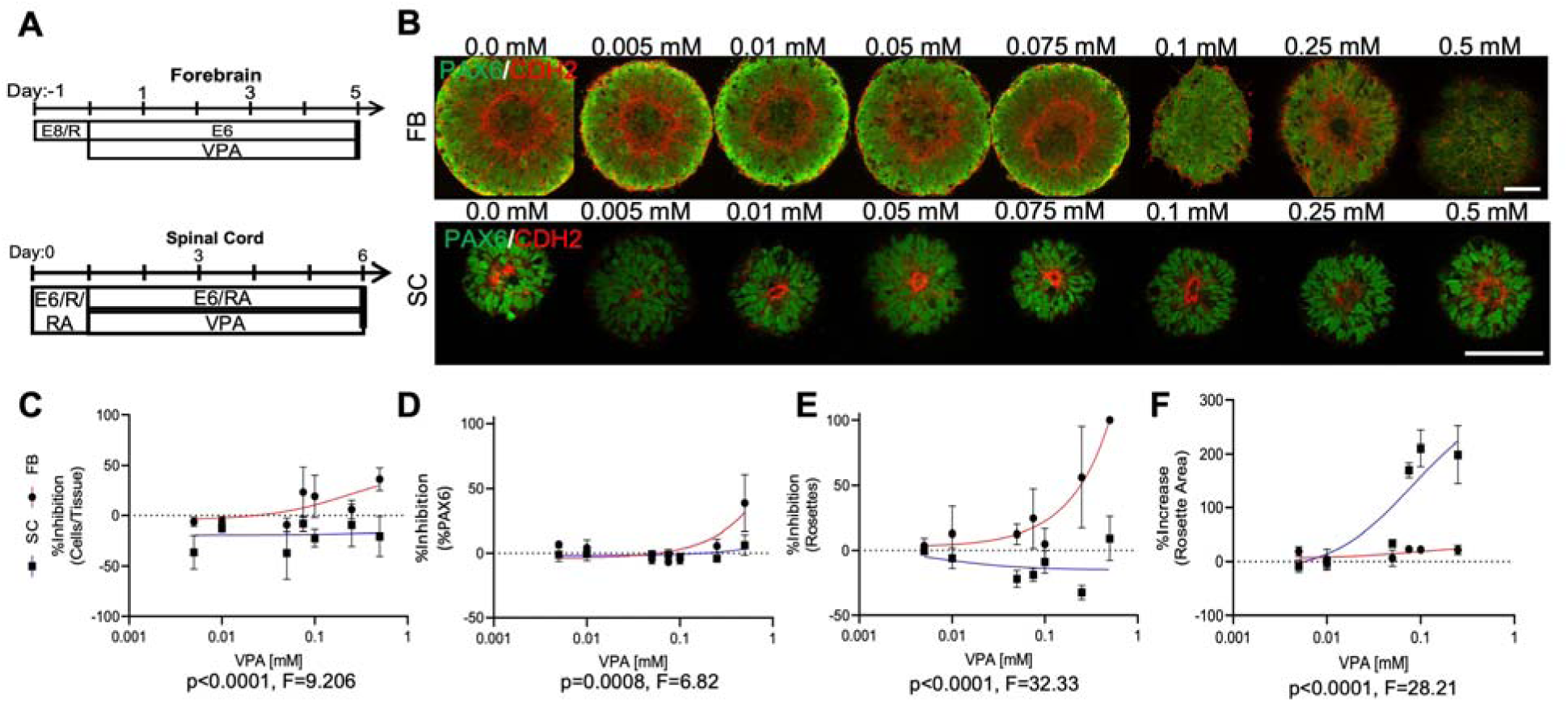
RosetteArrays display region-specific responses to VPA exposure. (A) Culture schema for FB & SC RosetteArray VPA dose-response experiments with (B) representative images of immunostained FB and SC rosettes. Comparison of FB and SC RosetteArray dose-response curves for %inhibition of (C) Cells/Tissue, (D) %PAX6 expression, and (E) single rosette emergence (Rosettes) as well as (F) measured N-cadherin^+^ polarized ring area. Each data point is the average of a WA09 hPSC (FB) or cervical spinal NMP progeny (SC) differentiation conducted in biological triplicate, n=50 technical replicates/well. Three parameter non-linear regression used to model and compare each metric. Scale bars are 100 µm.

### DNT screening using FB RosetteArrays with human metabolism integration

To evaluate the use of RosetteArrays for chemical DNT assessment, FB RosetteArrays were used to conduct a single concentration, proof-of-concept screen of 29 pharmaceutical and agrochemical compounds (**Figures 1A, 4**). The library contained 6 negative and 23 positive controls as determined by a literature review of general developmental toxicity or DNT, i.e., ‘Reference Literature Classification’ (**Figure S4**). Due to the 12-well plate format’s limited scalability, each compound was assayed at a single 10 μM concentration in (**Figures S5, S6, S8**). Thus, this screen was exploratory to evaluate the platform’s utility. Definitive DNT assessment should entail dose-response experiments with DNT control comparisons plus in vitro-to-in vivo extrapolation. For a single dose RosetteArray assay, the sample size for the ‘%Single Rosette’ metric is 3, i.e., the average of n=50 tissues analyzed per array/well, but the sample size for the ‘Cells/Tissue’, ‘%PAX6’, and ‘Rosette Area’ metrics is up to 150. Due to the statistical power of n=150, ‘Positive’ hit designation for these metrics required a statistically significance difference that was >25% decrease from control. If only a statistically significance decrease was observed, then it was considered a ‘Warning’. Additionally, for all compounds, the RosetteArray’s three primary metrics were assessed first, and if no positive hit was observed, then rosette area was also analyzed for further confirmation of non-toxic effects. Glycolic acid is the only exception to this rule given its well-known negative control status^43^. Lastly, to determine whether simulated human metabolism could be integrated into the RosetteArray platform, its tolerance for the presence of S9 pooled liver fraction was tested. Based on a dose-response, S9 is tolerated up to ∼0.002 mg/mL concentrations in the assay media solution (**Figure S7**). Seven pesticides were screened both with and without S9 pre-digestion for DNT.

**Figure 4.**
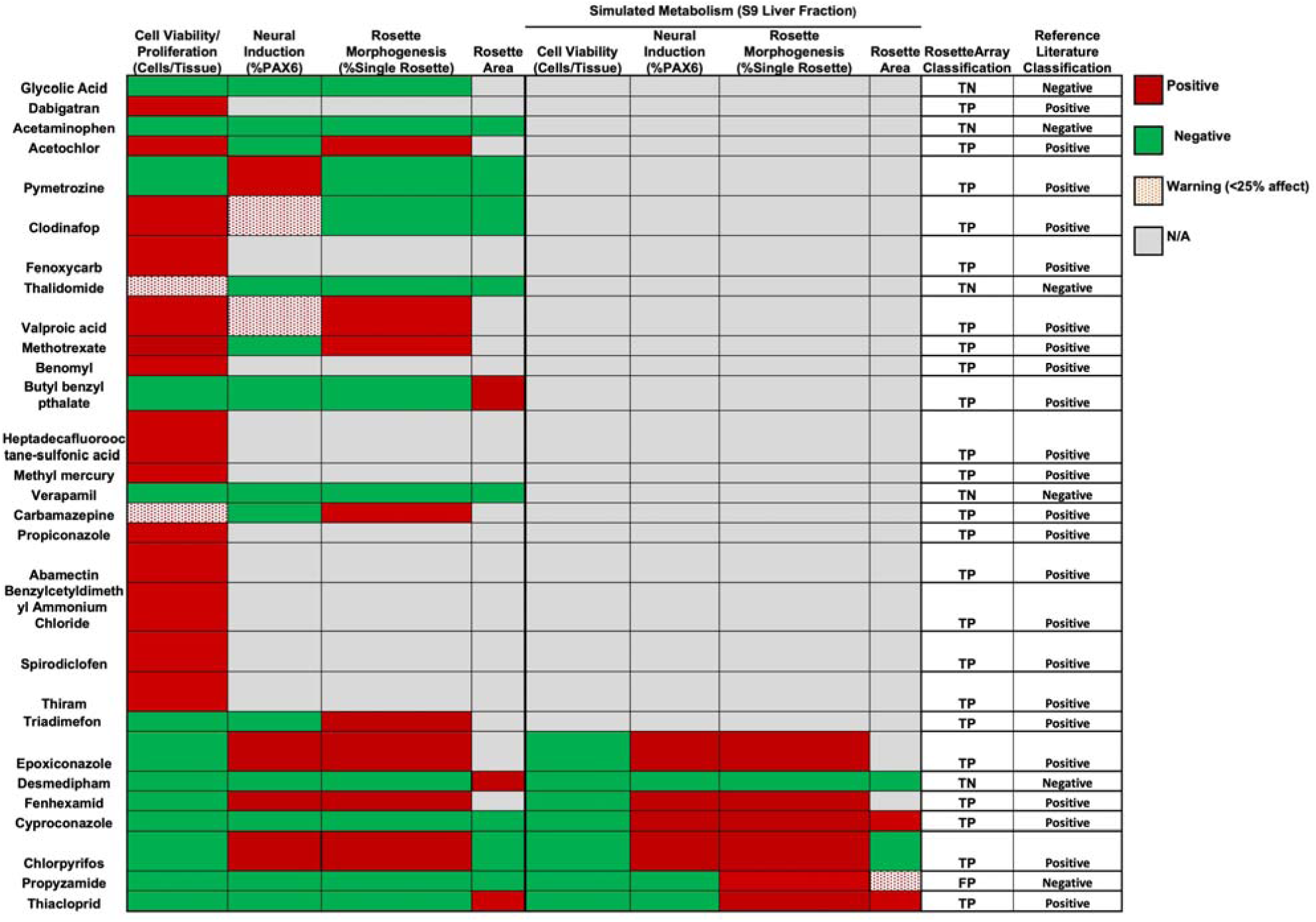
FB RosetteArrays can integrate simulated human metabolism and provide DNT assessment. RosetteArray classification for 29 select DNT compounds screened at 10 μM and with human S9 pooled-liver fraction pre-digestion for their ability to perturb cell viability/proliferation, neural differentiation, and rosette morphogenesis and area. Color coded summary: ‘Negative’, no effect - green; ‘Warning’, significant decrease >25% - dotted red; ‘Positive’ hit, significant decrease >25% - red; not analyzed – grey. Comparison of the FB RosetteArray’s ‘Assay Classification’ versus the compound’s ‘Reference Literature Classification’ is also provided. TN – true negative, TP – true positive, FN – false negative, FP – false positive. Non-metabolized compared to control via two-tailed T-test. Metabolized compared via one-way ANOVA. Each chemical was screened with a WA09 hPSC differentiation conducted in biological triplicate, n=50 technical replicates/well.

In the absence of simulated human metabolism, 4 of 6 negative control compounds tested as true negatives (**Figure 4**, ‘TN’). Desmedipham, an herbicide, was detected as ‘hit’ due to a statistically significant decrease in rosette area, but it is not generally considered to have DNT effects^54^. Also, thalidomide is a well-known teratogen; however, it was referenced as a DNT negative because its teratogenic effects on limb formation are due to inhibiting angiogenesis^55^. Our screen indicated it as a ‘warning’ due to inhibition of cell viability/proliferation. Of the positive controls, 22 out of 23 produced a hit with differential profiles indicating nuanced modes of action for observed DNTs. Cyproconazole is the only referenced positive control that did not produce a hit in the absence of simulated human metabolism. Of note, Butyl benzyl phthalate is a plasticizer with known reproductive toxicity^56^ and DNT in rodents^57^, but there is limited human data regarding its DNT affect. In this screen, it only significantly perturbed rosette area potentially indicating a subtle effect. Also, Chlorpyrifos was a clear hit in this initial screen, supporting recent efforts to bans its use as a pesticide^58,59^.

Seven pesticides were tested in the presence of simulated human metabolism (**Figure S8**). Desmedipham’s previous toxicity disappeared with S9-predigestion prior to screening. Alternatively, Cyproconazole and Propyzamide became toxic upon being metabolized (**Figure 4**). This correlates with Cyproconazole’s known disruption of Zebrafish development, including spinal malformations and locomotor defects^60,61^. Also, while Propyzamide has no known DNT effects, it is a known neurotoxicant in adult rats, alters gene expression associated with human developmental processes, and increases CYP450 enzymes upon human liver cell exposure^62^. Moreover, it disrupts tubulin-mediated microtubule assembly in plant cells^63^, a process observed in hPSC-derived rosette morphogenesis^20^. Thus, while a Propyzamide-induced DNT was not observed without simulated human metabolism in our assay or in other non-human model organism studies^61,64^, it shows clear DNT in the FB RosetteArray after being metabolized, placing doubt on its ‘Negative’ reference classification (**Figure 4, S8**). Across both metabolized and non-metabolized screens, the FB RosetteArray provided perfect specificity and near-perfect sensitivity DNT results, with the caveat of the studies small sample size and single-concentration evaluation. This demonstrates the platform’s ability to integrate simulated human metabolism and potential for human DNT risk assessment.

### Scale up to 96-well plate FB and SC RosetteArray assays

While effective, the 12-well plate RosetteArray format is insufficient for qHTS applications due to limited throughput and the amount of test compound required. For scale up to a 96-well plate format, Willow^®^ glass sheets were micropatterned and attached to bottomless 96-well plates using a double-sided adhesive as previously described^65^ (**Figure 5A**). FB RosetteArray scale up experiments demonstrated that the 12-well plate protocol did not directly translate to the 96-well format (**Figure 1C, 5B-E**). In both E6 and 0.1%DMSO supplemented media conditions, culture until Day 7 was required to reach acceptable single rosette emergence efficiencies (i.e., 76.06 ± 14.94% and 86.27 ± 9.06%, respectively). Using Day 7 as the termination timepoint, the optimized 8-day FB RosetteArray protocol was reproducible across full 96-well plate screens. When conducted in quadruplicate, with 0.1%DMSO supplemented media, and using only the interior 60-wells, plate-wide averages for single rosette emergence were 91.83 ± 5.80%, 95.46 ± 5.81%, 86.67 ± 6.84%, and 91.46 ± 5.81% (**Figure S9A**). Additionally, the FB RosetteArray’s Z-factor value is 0.532 across seventeen independent DNT screens using 1μM MTX and E6/0.1%DMSO as our positive and negative control conditions, respectively, indicating its suitability for qHTS (**Figure S9B**).

**Figure 5.**
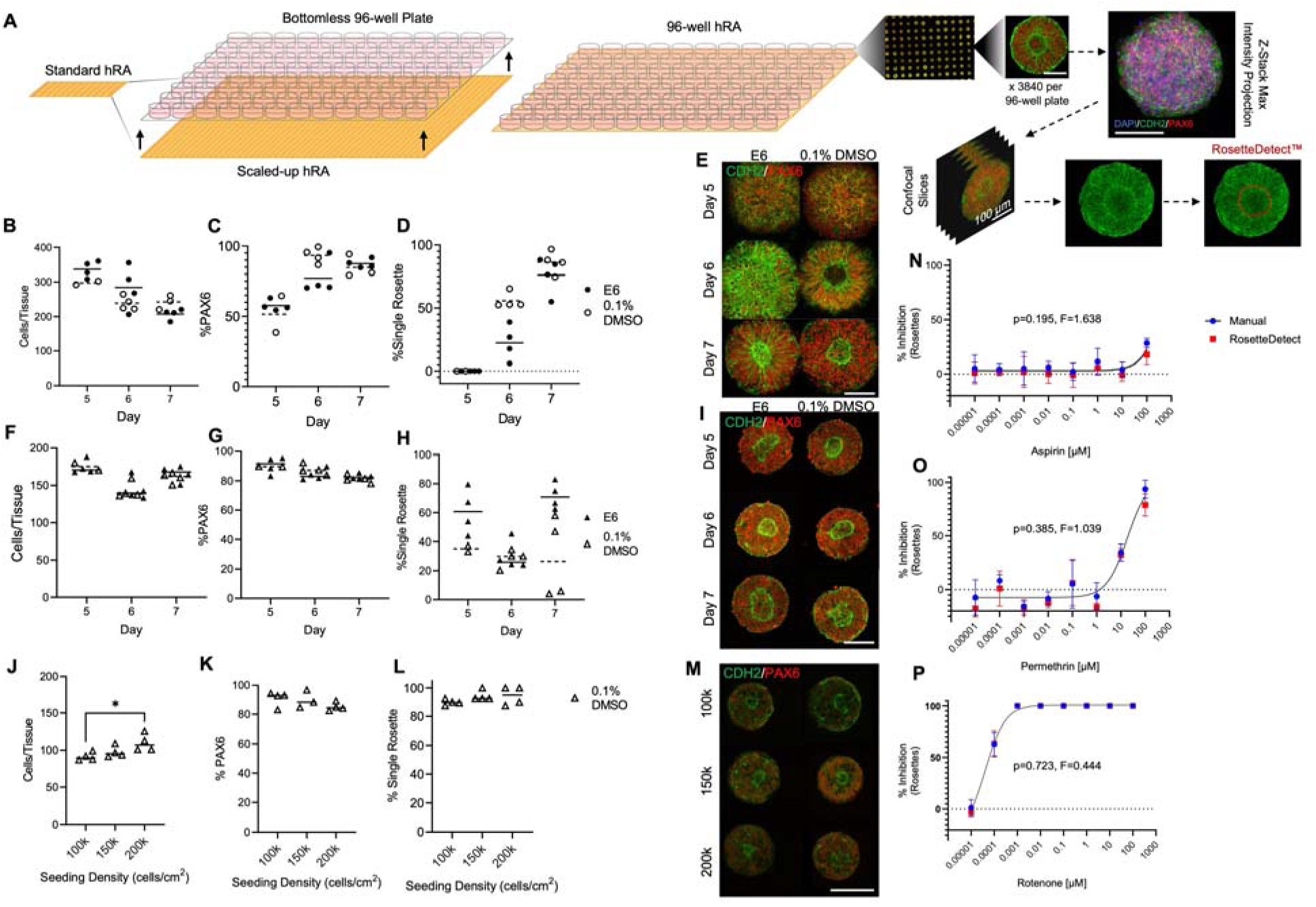
FB and SC RosetteArray assay and analysis scale-up to 96-well plate format. (A) Schematic of 96-well RosetteArray plate manufacture, image acquisition, and RosetteDetect image analysis. Quantification of (B-E) FB and (F-I) cervical spinal 96-well RosetteArrays with and without 0.1% DMSO across differentiation Days 5-7 for (B, F) cells/tissue, (C, G) neural induction, and (D, H) rosette emergence plus (E, I) representative immunostaining. Quantification of Day 5 lumbar spinal RosetteArrays with 0.1% DMSO at varied seeding density for (J) cells/tissue, (K) neural induction, and (L) rosette emergence plus (M) representative immunostaining. Each data point represents the average of a WA09 hPSC (FB) or cervical or lumbar spinal NMP progeny (SC) differentiation conducted in biological replicate with n=40 technical replicates per well. Dose-response curves for manually and RosetteDetect quantified FB RosetteArray assays of (N) Aspirin, (O) Permethrin, and (P) Rotenone. Each data point represents the average of a WA09 hPSC (FB) differentiation conducted in biological triplicate, n=40 technical replicates per well. Scale bars are 100 µm. Significance in dot plots assessed using 1-way ANOVA with Tukey-Kramer post-hoc analysis, * p ≤ 0.05. Dose-response curves compared using three-parameter non-linear regression models. If curves are statistically equivalent, then only one curve is displayed.

For cervical SC RosetteArrays, direct protocol translation yielded even worse single rosette emergence results (**Figure 5F-I**). For 0.1% DMSO media conditions, the 96-well plate cervical SC RosetteArray showed its best efficiencies at Day 5 of culture, but still much lower than the FB RosetteArray efficiency (**Figure 5D,H**). To improve the SC assay, its R/C regionalization and micropattern dimension were revisited in the 12-well format. Both cryopreserved cervical and lumbar-patterned NMPs^22^ were seeded onto micropatterns of 100, 125, and 150 μm diameter circles and assessed for single rosette emergence at Day 5 of culture (**Figure S10A**). While single rosette emergence within cervical tissues did not vary significantly across the micropattern dimensions, lumbar-patterned rosettes displayed increased (i.e., up to 93.80 ± 4.10%) single rosette emergence on 100 μm diameter micropatterns (**Figure S10B-I**). When the lumbar SC RosetteArray protocol was directly translated to the 96-well plate format, similar single rosette emergence efficiencies were observed in 0.1% DMSO supplemented media (**Figure 5J-M**).

RosetteArray analysis became a challenge with successful scale-up to 96-well plate formats that yielded ∼200 FB and ∼800 SC rosette tissues per well (**FigureS10J-L**). If automated confocal microscopy is used to acquire just 40 micropatterned tissues/well, then single rosette emergence within 3840 Z-stack images per plate would need to be analyzed manually (**Figure 5A**). Therefore, we utilized commercially available RosetteDetect™ software based on convolutional neural networks^66^ to automate rosette detection and segmentation within such images. The artificial intelligence (AI) model was trained on 588 images and tested on a separate 5594 image set from DNT dose-response studies. Model validation studies on ‘yes/no’ identification of single rosette emergence yielded accuracy, precision, recall/sensitivity, specificity, and F1 values of >90% (**Figure S9C,D**). Also, the model’s ability to segment the single rosette’s polarized N-cadherin ring structure was tested across 128 images producing a Dice Coefficient of 88.64 ± 6.50%, indicating good agreement of the rosette’s estimated area with manually curated ground truth data (**Figures 5A, S9E**). As a final demonstration of utility, RosetteDetect’s performance in quantifying single rosette emergence across 3 DNT RosetteArray assays, i.e., Aspirin, Permethrin, and Rotenone, was compared to manual analysis. Dose-response curves generated using either manual or RosetteDetect analyses were statistically equivalent (**Figures 5N-P, S9F**). With inclusion of CellProfiler™ coding to quantify DAPI^+^ and PAX6^+^ cells/image slice, these results collectively demonstrate feasible scaling of RosetteArray assays and RosetteDetect analysis for qHTS applications.

### RosetteArrays detect chemical NTD risks specific to folate metabolic pathway inhibition

Of NTD risk pathways surveyed in Figure 2, disruption of folic acid metabolism is the best known NTD risk factor^48^. Therefore, the RosetteArray platform’s ability to detect folate metabolic pathway-specific NTD risk was interrogated further. First, we assessed the ability to rescue MTX inhibition of rosette emergence by supplementing with the metabolized form of folic acid, 5-methyltetrahydrafolic acid (5-MTHFA)^67^. 12-well FB RosetteArrays were used to conduct a MTX dose-response in the absence and presence of 20 μM 5-MTHFA (**Figures 6A-F, S11A, B**). In all metrics, the presence of 20 μM 5-MTHFA significantly shifted the dose-response curve to the right increasing IC_50_ values by greater >10-fold. This demonstrates folate metabolic pathway-specific rescue of MTX’s inhibitory effects.

**Figure 6.**
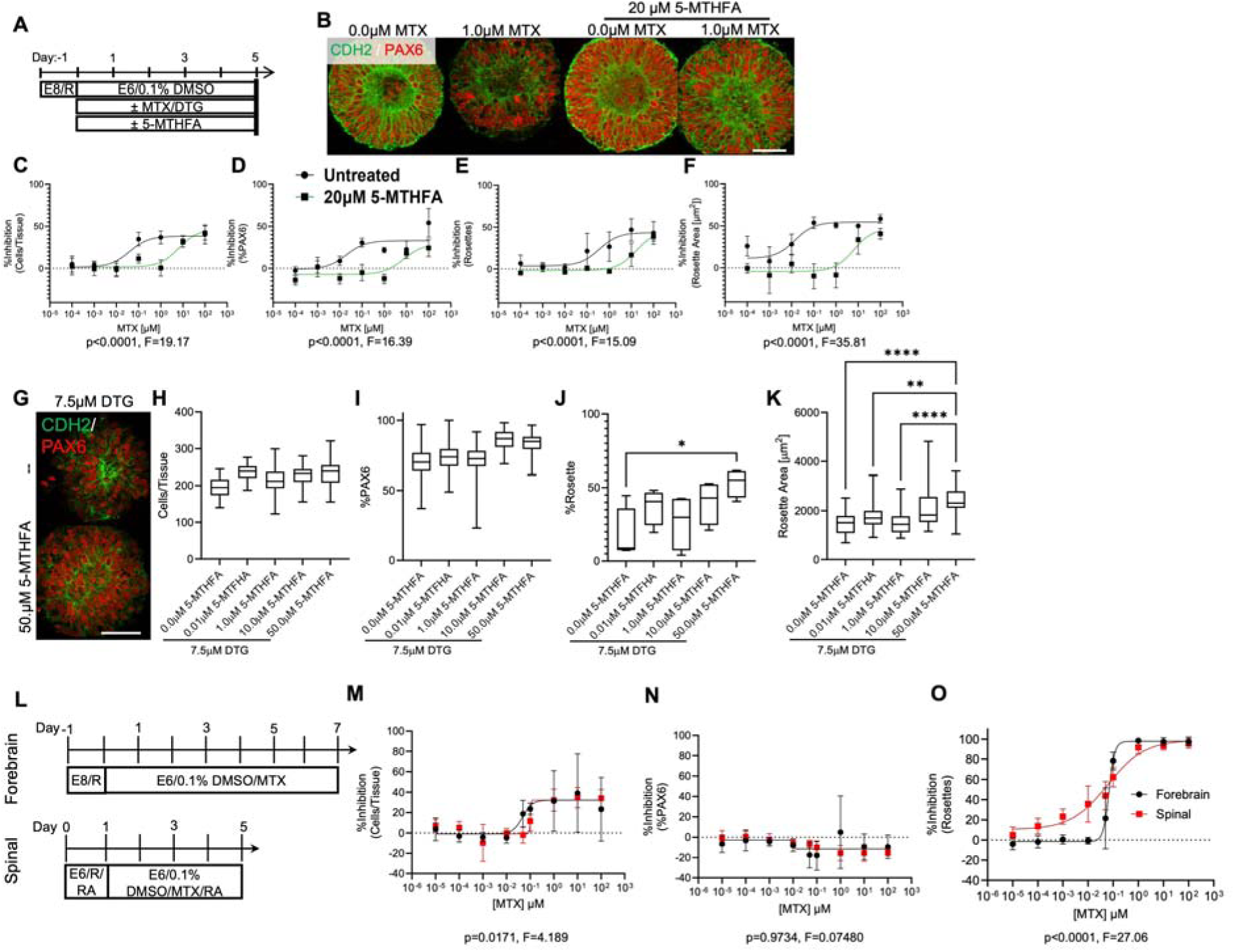
FB and SC RosetteArrays detect folate metabolic pathway-specific risk with regional differences. (A) Culture schema for MTX/DTG exposure with 5-MTHFA rescue in 12-well FB RosetteArrays (WA09 hPSC seeded). (B) Inhibitory effect of 1.0 µM MTX on FB rosette emergence and rescue using concurrent 20.0 µM 5-MTHFA treatment. Quantification of MTX dose-response on 12-well FB RosetteArray derivation with 20.0 µM 5-MTHFA treatment across (C) cells/tissue, (D) neural induction, (E) single rosette emergence, and (F) rosette area metrics. Quantification of DTG’s inhibitory effects on 12-well FB RosetteArray derivation with increasing 5-MTHFA supplementation across (H) cells/tissue, (I), neural induction, (J) single rosette emergence, and (K) rosette area metrics. (L) Culture schema for MTX dose-response comparison between 96-well FB and lumbar SC RosetteArrays. Dose-response curves of %inhibition of (M) cells/tissue (N) neural induction, and (O) single rosette emergence. (C-F, H-K) Each data point is the average of a WA09 hPSC (FB) differentiation conducted in biological triplicate or quadruplicate, n=50 technical replicates per well. (C-F) Three-parameter non-linear regression used to model and compare each metric. (H-K) Significance assessed using 1-way ANOVA with Tukey-Kramer post-hoc analysis, * p ≤ 0.05, **p ≤ 0.01, ***p ≤ 0.001, ****p ≤ 0.0001. (M-O) Each data point is the average of WA09 hPSC or lumbar spinal NMP progeny differentiations each conducted in biological octuplet, n=40 technical replicates per well. Four-parameter non-linear regression used to model and compare each metric. For (N), curves are statistically equivalent, thus only one curve is displayed. Scale bars are 100 µm.

Second, the 12-well FB RosetteArray was used to detect the NTD risk posed by Dolutegravir, an HIV antiretroviral therapy. When taken periconceptionally, Dolutegravir has been shown to increase clinical NTD occurrence^68,69^, presumably by antagonistically binding the folate receptor-1^67^. Moreover, its NTD risk effects can be negated by sufficient folate dietary supplementation^67,70^. In our FB RosetteArray dose-response, Dolutegravir inhibits all metrics similarly with a single rosette emergence IC_50_ of 3.19μM (2.15-4.17μM, 95% CI), which is within its therapeutic serum level of 3-10 μM^67^ (**Figure S11C-G**). This is indicative of its presumed inhibition of folate receptor-1, which would inhibit vital folate metabolism required for basic cell physiology. Additionally, analogous to MTX, conducting the FB RosetteArray at 7.5 μM DTG with increasing 5-MTHFA concentrations showed a significant rescue in single rosette emergence and rosette area with non-significant but increasing trends in cell viability/proliferation and neural induction metrics (**Figure 6G-K, S11H**). Collectively, these results demonstrate that the FB RosetteArray can detect folate metabolic pathway-specific NTD risk associated with both MTX’s inhibition of the pathway’s dihydrofolate reductase enzyme and DTG’s inhibition of folate receptor-1 binding^67^.

Third, due to previous regional differences in RosetteArray responses to VPA (**Figure 3**), FB and lumbar SC RosetteArray detection of MTX’s NTD risk was compared using the 96-well plate format (**Figure 6L-O**). Relative to the 12-well assay, the 96-well FB RosetteArray showed increased sensitivity to MTX’s inhibitory effects with a >5-fold decrease in the single rosette emergence IC_50_ value, i.e., 0.069 μM (0.062-0.076 μM, 95% CI) vs. 1.4 μM (0.61-3.83 μM, 95% CI) vs. 0.351 μM (0.3-2.42 μM, 95% CI) (**Figures 6O, 2O, 6E, respectively**). Also, maximal inhibition of single rosette emergence was observed at ∼1 μM vs. >100 μM in the 96- vs. 12-well FB RosetteArray formats (**Figures 6O, 2O**). When evaluating the 96-well plate format’s regional differences, the lumbar SC RosetteArray had a statically equivalent IC_50,_ i.e., 0.61 μM (0.035-0.10 μM, 95% CI), to the FB RosetteArray. Yet, its Lowest Observed Adverse Effect Level (LOAEL) was 500-fold lower than the FB RosetteArray, i.e., 0.1 nM (SC) vs 0.5 μM (FB) (**Figure 6O**). Moreover, above the IC_50_, MTX inhibited single rosette emergence in SC and FB RosetteArrays similarly, but below the IC_50_, SC tissues appeared more prone to form 2+ versus no rosettes per tissue (**Figures S12A-C**). Again, this highlights regional differences between rosette emergence mechanisms in FB vs. SC RosetteArrays mirroring presumed differences between regional NTD mechanisms *in vivo*^71^.

### RosetteArrays detect genetic and multifactorial NTD risks

In addition to chemical exposure risks, genetic variants of folate metabolic and planar cell polarity (PCP) pathway machinery are also known to increase NTD risk^12^. To evaluate the RosetteArray platform’s ability to detect such genetic risk factors, the WA09 hESC line was gene edited to generate clones with clinical *MTHFR^C677T^*(p.A222V)^72^ and *SCRIB^C3128T^* (p.P1043L)^73^ variants. In the folate metabolic pathway, methyltetrahydrofolate reductase (MTHFR) generates 5-methyltetrahydrofolate (5-MTHFA), which is the metabolized folate form found in blood plasma^72^. The c.C677T mutation decreases the enzyme’s activity by ∼35% in heterozygotes and ∼70% in homozygotes^74^. Sequencing revealed that the WA09 hESC line was heterozygous for the c.C677T mutation. Therefore, *MTHFR^C677T(-/-)^* homozygote clones #4 and #9 were generated using gene editing (**Figure S13A,B**). In the PCP pathway, the Scribble protein (SCRIB) acts to scaffold protein-complexes at the plasma membrane during epithelial cell apical-basal polarization^75^. Its N-terminal Leucine-Rich-Region (LRR) enables membrane localization, while its four PDZ domains, with the c.C3128T/p.P1043L mutation in its third PDZ domain, are reported to also effect membrane localization and PCP protein interactions^73,75^ (**Figure S13C**). Gene editing of the WA09 parent line yielded a heterozygous (SCRIB^P1043L(+/-)^, #78), a homozygous (SCRIB^P1043L(-/-)^,#52), and a knockout (SCRIB^KO^, #17) clone. The knockout was caused by a downstream deletion yielding a nonsense frameshift (**Figure S13D-G**). All clones could generate rosette containing cultures in standard 6-well forebrain NEC derivation experiments^29^ (**Figure S13H-J**). Also, the parent and isogenic mutant lines had normal karyotypes with no mycoplasma detection (**Figure S14**).

To begin assessing the RosetteArray’s sensitivity to NTD risk caused by the *MTHFR^C677T(-/-)^* variant, the *MTHFR^C677T(+/-)^* WA09 parent line and homozygous clones #4 and #9 were screened using 12-well FB RosetteArrays. The assay’s media contains 9 μM folic acid. Therefore, this initial screen was run using control, 1 μM MTX, and 1 μM MTX plus 20 μM 5-MTHFA media conditions (**Figure S15A,B**). Of note, this experiment simulates a multifactorial NTD risk scenario, where the combination of the *MTHFR^C677T^* genetic variant and a MTX exposure are being modeled. No difference between lines in any metric is observed under control FB RosetteArray conditions (**Figure S15C-E**). However, upon challenging with 1 μM MTX, all lines showed a statistically significant decrease in cell viability/proliferation and single rosette emergence metrics. Yet, both *MTHFR^C677T(-/-)^* clones demonstrated significantly increased MTX sensitivity in single rosette emergence, i.e., 13.62 ± 11.27% (Clone #4) and 23.47 ± 6.81% (Clone #9), compared to the *MTHFR^C677T(+/-)^* parent line, i.e., 51.50 ± 1.06%. This effect is folate metabolic pathway specific since 20 μM 5-MTHFA supplementation, in the presence of 1 μM MTX, rescued the risk phenotype in all lines.

To further clarify differences in folate metabolic pathway sensitivities, a full MTX dose-response in a 96-well FB RosetteArray was conducted comparing the *MTHFR^C677T(+/-)^* parent line, *MTHFR^C677T(-/-)^*clone #9, and the *MTHFR^C677T(+/-)^/* SCRIB^P1043L(-/-)^ clone (**Figures 7A,B, 15F-I**). Each lines’ dose-response yielded statistically different curves with the *MTHFR^C677T(-/-)^* clone #9’s IC_50_ of 0.0305 μM (0.0200-0.0464 μM 95% CI) being lower than the *MTHFR^C677T(+/-)^/* SCRIB^P1043L(-/-)^ clone’s and *MTHFR^C677T(+/-)^* parent line’s IC_50_s of 0.0411 μM (0.0256-0.0656 μM 95% CI) and 0.0477 μM (0.0327-0.0686 μM 95% CI), respectively. For increased accuracy, a second MTX dose-response between the *MTHFR^C677T(-/-)^* clone #9 and its *MTHFR^C677T(+/-)^* parent line was conducted in duplicate with additional dosages around the previously observed IC_50_ values (**Figure 7C, S15J-M**). Again, each lines’ dose-response yielded statistically different curves but now without overlapping IC_50_ values, i.e., Clone #9- 0.0497 μM (0.0452-0.0544 μM 95% CI) vs. WA09 parent- 0.0697 μM (0.0629-0.0768 μM 95% CI). Additionally, a 5-MTHFA rescue dose-response in the presence of 1 μM MTX showed that *MTHFR^C677T(-/-)^*clone #9 required more folate supplementation, i.e., EC_50_ of 7.90 μM (5.38 −11.20 μM 95% CI), than the *MTHFR^C677T(+/-)^* parent line, i.e., EC_50_ of 3.70 μM (2.57 −5.20 μM 95% CI) to rescue its risk phenotype (**Figure 7D-F, S15N-P**).Thus, the *MTHFR^C677T(-/-)^* lines’ increased sensitivity to folate metabolic pathway perturbation is due to its additional c.C677T mutation compared to the *MTHFR^C677T(+/-)^* parent line. Moreover, this highlights the RosetteArray platform’s ability to quantify a genetic NTD risk factor in a multifactorial scenario.

**Figure 7.**
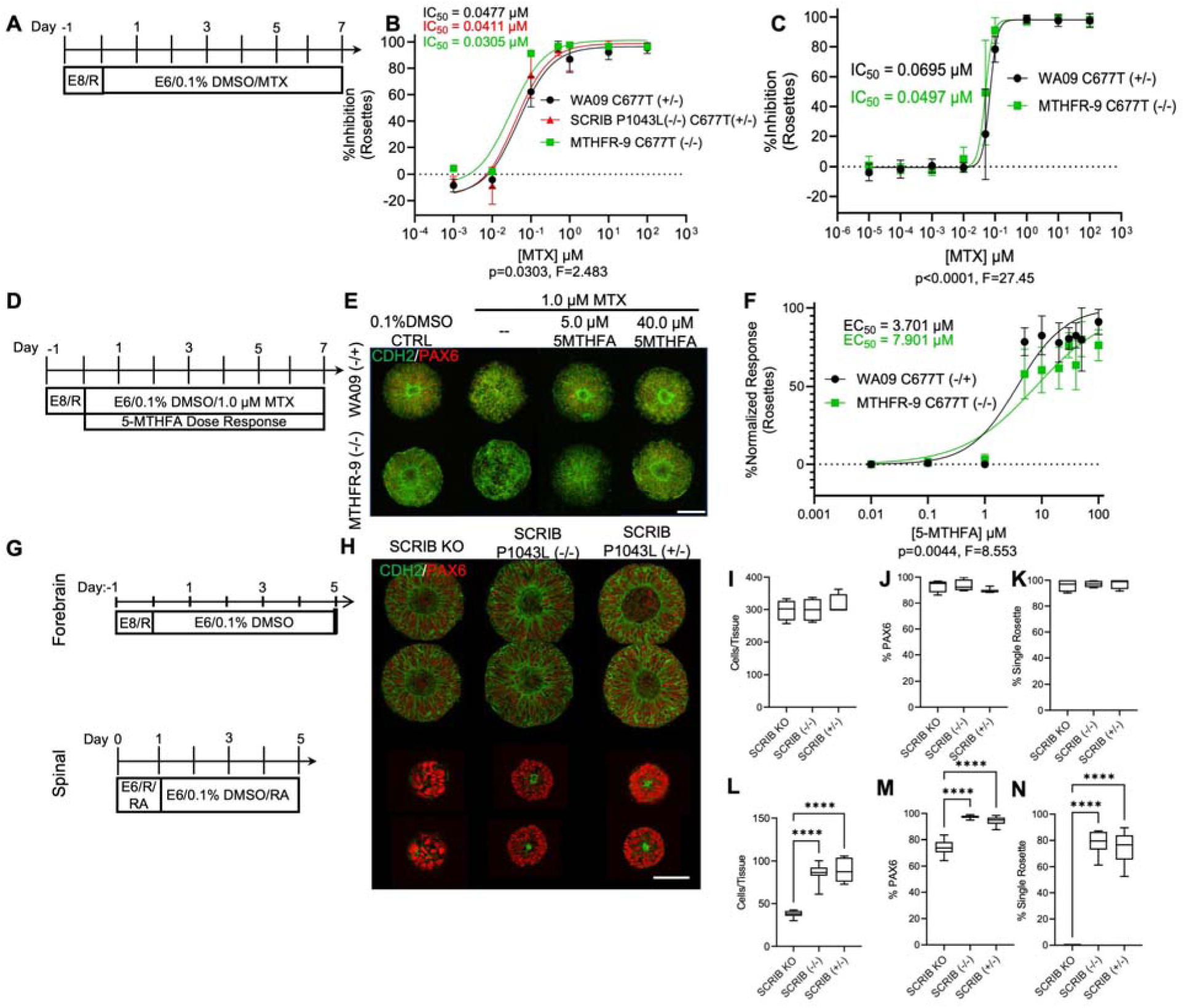
RosetteArrays detect clinically relevant genetic and multifactorial NTD risks. (A) Culture schema for (B) initial and (C) secondary FB RosetteArray MTX dose-response comparison of single rosette emergence across *MTHFR^C677T(-/-)^* and SCRIB^P1043L(-/-)^/*MTHFR^C677T(+/-)^* mutant lines and the WA09 *MTHFR^C677T(+/-)^* isogenic control. (D) Culture schema for 5-MTHFA dose-response rescue of *MTHFR^C677T(-/-)^* mutant and the WA09 *MTHFR^C677T(+/-)^* isogenic control under 1 μM MTX exposure with (E) representative staining and (F) quantification of %normalized single rosette emergence. Each data point is the average of a WA09 or mutant hPSC differentiation conducted in biological quadruplicate, n=40 technical replicates per well, except for (C), which is the average of two WA09 and mutant hPSC differentiations. Four-parameter non-linear regression used to model and compare each metric. (G) Culture schema for FB and cervical SC RosetteArrays of SCRIB^P1043L(+/-)^, SCRIB^P1043L(-/-)^, SCRIB^KO^ mutant lines with (H) representative immunostaining and FB (I-K) and SC (L-N) quantification for cells viability/proliferation, neural induction, and %single rosette emergence. Data represents (I-K) a hPSC differentiation conducted in 4 biological replicates (n=40 technical replicates per well) or (L-N) two cervical spinal NMP differentiations each conducted in n=3-4 biological replicates (n=40 technical replicates per well). Data is compared via a 1-way ANOVA with Tukey-Kramer post-hoc analysis. ****p ≤ 0.0001. Scale bars are 100 µm.

To model PCP pathway-induced NTD risk, SCRIB^P1043L(+/-)^, SCRIB^P1043L(-/-)^, and SCRIB^KO^ lines were generated and cryopreserved as both hESC and cervical spinal NMP banks along with the WA09 parent line (**Figure S16A,B**). In 12-well FB RosetteArrays, no difference was observed between the 3 mutant lines despite immunostaining showing FB rosette formation in the absence of SCRIB protein for the knockout mutant (**Figures 7G-K, S16C**). Yet, in the 12-well cervical SC RosetteArray, there was complete abrogation of rosette emergence accompanied by a significant decrease in cell viability/proliferation and neural induction in the SCRIB^KO^ line compared to the SCRIB^P1043L(+/-)^ and SCRIB^P1043L(-/-)^ lines (**Figure 7G,H,L-N**). This regional NTD risk phenomenon in the SCRIB^KO^ line comports with the *SCRIB^Crc^* mouse^75^ and human^76^ presentation of craniorachischisis (CRN), which is a failure of neural tube closure across the neuraxis except for the rostral forebrain^77^. Also, despite SCRIB’s p.P1043L mutation in its third PDZ domain, immunostaining showed apical co-localization of SCRIB and F-Actin (phalloidin) in both FB and cervical SC rosettes formed using SCRIB^P1043L(+/-)^ and SCRIB^P1043L(-/-)^ lines (**Figure S16C**). However, SCRIB^KO^ rosettes showed F-Actin apical polarization in FB Rosettes despite minimal-to-no SCRIB immunostaining, and no F-actin polarization in cervical SC rosettes. In mice, others have observed that SCRIB mutations, which lower expression (*SCRIB^rumz/rumz^*) causing CRN, also cause decreased apical immunostaining of F-actin and other tight junction proteins in defect presenting regions^75^.

This striking R/C regional difference led us to investigate expression of SCRIB in FB and cervical SC rosette tissues. In humans, SCRIB has two isoforms (A and B) with SCRIB A containing a c-terminal AbLIM domain that links the protein to F-actin through binding β spectrins^78^ (**Figure S13C**). Using qPCR for all SCRIB and its isoforms, we observed a decreasing trend in expression of all SCRIB and isoform B as cells transitioned from hESC to FB NEC or to SC NMP and NEC states (**Figure S16D**). Yet, only the WA09 parent lines’ SCRIB B decrease during this transition was statistically significant. However, the SCRIB A isoform’s expression trend slightly increases in FB NECs in all lines except for the SCRIB^KO^, confirming its knockout status. Overall, the composite data suggests a FB-SC regional difference in the importance of SCRIB for NEC apical-basal polarization that motivates further investigation. Additionally, it again highlights potential R/C regional differences in NTD mechanisms and motivates NTD risk assessment across the neuraxis as enabled by the RosetteArray platform.

## DISCUSSION

The RosetteArray platform standardizes morphogenesis of hPSCs into incipient forebrain and spinal neural organoids, a.k.a. single rosette tissues^21^, thereby providing a predictable analogue of human neural tube emergence for screening applications. To our knowledge, it’s the first standardized hPSC-derived rosette/neural tube morphogenesis assay that can be generated using direct seeding of cryopreserved cells, thereby enabling an off-the-shelf assay^24,25,27,30^ (**Figure 1**). Our results demonstrate its potential as a human neurodevelopmental hazard assessment tool for general DNT screening and for modeling NTD risk factors. Evaluation with 31 different chemicals with known or presumed NTD risk/DNT profiles demonstrated that the RosetteArray yields preliminary DNT specificity and sensitivity values of 1.0 and 0.83, respectively (**Figures 2, 4, S11C-G**). The only negative control missed was Propyzamide, of which its ‘negative’ assignment in a human metabolized scenario is questionable^62,63^. Also, our experiments showed that the platform, using both FB and SC RosetteArrays, can reliably detect environmental/chemical, genetic, and multifactorial NTD risk factors (**Figures 5, 6**). Interestingly, some NTD risk factors demonstrated region-specific perturbations of rosette morphogenesis (**Figures 3, 7G-N**), indicating the importance of encompassing diverse neuraxial regions in DNT assays. Lastly, with scaling to a reproducible (Z-factor = 0.532) 96-well plate format plus AI-based image analysis (**Figures 5, S10**), the RosetteArray could enable qHTS of broad human neurodevelopmental risk as demonstrated here for NTDs and elsewhere for Autism Spectrum Disorder (ASD)^20^. As shown in the ASD exemplar, it is important to note that a RosetteArray ‘hit’ does not mean that rosette morphogenesis cannot occur, it only means that the morphogenesis was perturbed enough to detect a deviation from the norm. The RosetteArray platform makes such detection drastically more scalable and thereby sensitive.

Nearly two decades ago, US regulatory agencies stated the need for developing New Alternative/Approach Methodologies (NAMs) to better recapitulate human biology for risk assessment purposes and to reduce the use of animal testing^79,80^. Specific to DNT, the Organization for Economic Cooperation and Development recently released a DNT in-vitro testing battery guideline, which includes assays for neural progenitor cell proliferation, apoptosis, and cell migration but lacked a scalable neural rosette formation assay^81^. The RosetteArray platform provides this assay by enabling spatial and temporal prediction of rosette morphogenesis, which encompasses hPSC viability, proliferation, neural differentiation, and NEC viability, proliferation, and rosette morphogenesis^34^. Additionally, we and others have used rosette morphogenesis to assess risk of other neurodevelopmental^16,20^ and neurodegenerative^26,33^ disorders. Thus, further exploration into the extent that inclusion of the RosetteArray assay could streamline the DNT in-vitro testing battery is warranted. This would help achieve the regulatory goal of providing optimal human DNT risk assessment while balancing industry’s goal for efficient and cost-effective tools for safety assessment in discovery pipelines. Of note, chemical exposure during FB RosetteArray assays occurs throughout the hPSC (Day 0) to NEC rosette formation (Day 5 or 7) process, thereby enabling possible detection of general developmental toxicity in addition to specific DNT (**Figure 6A, 7A**).

NTDs are well known to have environmental, polygenic, and multifactorial etiologies, which have historically been investigated using mouse models^12^. This approach has and continues to provide fundamental insights. Yet, it has inherent translational and experimental limitations. Translational limitations caused by differences between mice and humans include genomic sequence/architecture, number of neural tube closure points, and the fact that mouse models predominantly display cranial defects versus spina bifida defects, which is the majority of human clinical cases^71,82^. Experimentally, the limited number of mouse strains used to create NTD models impedes exploration of how population variation affects polygenic etiology^71^. Also, experimental possibilities are limited by the sheer number of mice required for regulatory DNT studies, i.e. n=20 per experimental group^83^, or to demonstrate the rate of NTD occurrence. For example, Tukeman et al. required the use of 169 dams/1302 fetuses to observe 7 NTD cases and demonstrate the folate-responsive NTD risk associated with periconceptional DTG administration^70^.

The RosetteArray platform helps to address these limitations by providing a scalable hPSC-derived model of FB and SC neural tube emergence. It displays regional differences in morphogenesis^21^ (**Figures 1D, 5E,I,M, and S10**) and NTD risk responses (**Figure 3, 6O, 7G-N, S12**) that may be indicative of differences in neural tube closure mechanisms^71^. Using FB RosetteArrays, DTG’s folate-responsive NTD risk was detected at therapeutic serum levels (**Figure 6G-K, S11C-G**), further supporting clinical^69^ and rodent studies^70^. Also, since the platform can be derived using iPSC lines^20^, it could enable population variation studies to, for example, elucidate polygenic and multifactorial etiologies underlying *MTHFR^C677T^* penetrance in NTD cases. Using just two FB RosetteArray 96-well plates, we observed an increased NTD risk (TC vs. TT IC_50_ ratio = 1.40) associated with *MTHFR^C677T^* homo- vs. heterozygosity in the WA09 genetic background (**Figure 7C**). A finding recapitulated in meta-analysis of 19 different clinical studies with ∼6438 human participants that yielded a TC vs. TT odds ratio of 1.427 (1.247-1.634, 95% CI)^84^. While IC_50_ ratios and odds ratios are not equivalent, it does support further use of the RosetteArray for conducting NTD etiology studies to investigate population variance. Coupled with the platform’s ability to enable *in vitro* observation of SCRIB^KO^’s R/C region-specific NTD risk phenomena, which is clinically associated with CRN^75^, RosetteArray technology could have a transformative impact on investigating human NTD etiology, despite lacking non-neural components that surround the emergent neural tube *in vivo* (**Figure 2Q-T, S3F,J**)^25,85^. Moreover, the RosetteArray platform’s scalability could be applied to elucidate the etiology of other NDDs and enable a transformative level of qHTS of precision medicine therapeutics to prevent or mitigate such disorders.

## AUTHOR CONTRIBUTIONS

B.F.L, G.T.K., and R.S.A. took part in conception and design of all RosetteArray experiments and jointly interpreted all data. K.K. and R.W. conceived and designed all RosetteDetect experiments and jointly interpreted all related data. B.F.L. and G.T.K. performed all RosetteArray experimentation unless otherwise noted. N.J.K. performed all RosetteArray assays whose control data was included in Z-factor analysis. N.R.I. aided in derivation of WA09 and *SCRIB* mutant cell banks. F.S. performed long-term RosetteArray culture experiments and analysis. B.F.L., G.T.K., N.J.F., J.E.M., and M.R.C. performed data acquisition and data analysis. N.R.I. and J.E.M. assisted with platform manufacturing, immunostaining, and imaging. J.F.R. and B.J.I. assisted with design and interpretation of DNT and NTD experiments. B.F.L, G.T.K., F.S., and RSA drafted the manuscript for publication with comments from all authors.

## ACKNOWLEDGEMNTS

We thank the University of Wisconsin-Madison’s Human Pluripotent Stem Cell and Gene Editing Service for generating *MTHFR* and *SCRIB* mutant lines. They are supported, in part, by a NICHD U54HD090256 grant and UW2020 Grant awarded to Anita Bhattacharyya and Su-Chun Zhang by the UW-Madison and the Wisconsin Alumni Research Foundation. B.F.L, N.J.F., F.S., and R.S.A were supported by NICDH R21HD103111 and NSF ERC #1648035 grant awarded to the Ashton lab at UW-Madison. G.T.K., N.J.F., K.K., R.W., and R.S.A. were supported by NIEHS R42/43ES033912 grant awarded to Neurosetta LLC. B.F.L. was also supported by NIH MSTP T32GM14095. G.T.K. and R.S.A. were also supported by Draper TIF grant #AAH6969 and an Accelerator grant #AAI6923 awarded by WARF. N.R.I. and R.S.A. were also supported by U.S.LJEnvironmental Protection Agency grant 83573701. R.S.A. was also supported by NCATS UG3TR003150, NSF CAREER Award #1651645, and an Innovation in Regulatory Science Award #1014150 from the Burroughs Wellcome Fund. R.W. was also supported by NSF awards DMS-AWD00000326 and PHY2317138 and the Simons Foundation award MP-TMPS-00005320.

## STAR□Methods

### KEY RESOURCES TABLE

| REAGENT or RESOURCE | SOURCE | IDENTIFIER |
| --- | --- | --- |
| Antibodies |  |  |
| SCRIB primary | Invitrogen | (Thermo Fisher Scientific Cat# PA5-54821, RRID:AB_2647030) |
| N-cadherin primary | BD Biosciences | (BD Biosciences Cat# 610920, |
|  |  | RRID:AB_2077527) |
| PAX6 primary | BioLegend | (BioLegend Cat# 901301 (also 901302), RRID:AB_2565003) |
| β-tubulin primary | Biolegend | (Cat# 802001, RRID:AB_2564645 |
| Brachyury primary | R&D | Cat# AF2085, RRID:AB_2200235 |
| Sox2 primary | Millipore | Cat# MAB4423, RRID:AB_11213224 |
| Donkey anti-Rabbit IgG 488 | Invitrogen | (Thermo Fisher Scientific Cat# A-21206 (also A21206), RRID:AB_2535792) |
| Donkey anti-Mouse IgG 488 | Invitrogen | (Thermo Fisher Scientific Cat# A-21202, RRID:AB_141607) |
| 555 Donkey anti-Rabbit | Invitrogen | (Thermo Fisher Scientific Cat# A-31572 (also A31572), RRID:AB_162543) |
| Donkey anti-Mouse IgG 555 | Invitrogen | (Thermo Fisher Scientific Cat# A-31570, RRID:AB_2536180) |
| horseradish peroxidase-conjugated goat anti-rabbit | Jackson ImmunoResearch Laboratories | (Jackson ImmunoResearch Labs Cat# 111-035-003, RRID:AB_2313567) |
| Chemicals, Peptides, and Recombinant Proteins |  |  |
| ROCK Inhibitor (Y27632) | Tocris | Cat#: 1254 |
| FGF8b | PeproTech | Cat#: 100-25 |
| CHIR99021 | Tocris | Cat#: 4423 |
| GDF11 | PeproTech | Catalog Number:120-11 |
| Dorsomorphin | Tocris | Cat#: 3093/10 |
| Retinoic Acid | Sigma-Aldrich | Cat#: R2625 |
| Penicillin-Streptomycin | Invitrogen | Cat#: 15140122 |
| TRIzol | Invitrogen | Cat#: 15596026 |
| Triton X-100 | Sigma-Aldrich | Cat#:A16046.AE |
| DAPI | Invitrogen | Cat#: D1306 |
| Alexa Fluor 488 Phalloidin | Invitrogen | Cat#: A12379 |
| ω-mercaptoundecyl bromoizobutyrate | ProChimia | Cat#: FT 015-m11-0.2 |
| PEGMEMA | Sigma-Aldrich | Cat#: 447935 |
| sodium ascorbate | Sigma-Aldrich | Cat#: 1613509 |
| 2,2'-Bipyridine | Sigma-Aldrich | Cat#: D216305 |
| Copper (II) Bromide | Sigma-Aldrich | Cat#: 221775 |
| human S9 liver fraction | Sigma-Aldrich | Cat#: S2442 |
| CloneR | StemCell Technologies | Cat#: 05889 |
| TrueCut Cas9 Protein V2 | Thermo Fisher Scientific | Cat#: A36498 |
| QuickExtract DNA Extraction Solution 1.0 | Thermo-Fisher | Cat#: NC9904870 |
| protease inhibitors | Thermo-Fisher | Cat#: 78429 |
| Glycolic Acid | Sigma-Aldrich | Cat#: 124737 |
| Benomyl | Sigma-Aldrich | Cat#: 45339 |
| Valproic Acid Sodium Salt | Sigma-Aldrich | Cat#: P4543 |
| Methotrexate | Sigma-Aldrich | Cat#: 1414003 |
| Novobiocin | Sigma-Aldrich | Cat#: N6160 |
| Dabigatran | Sigma-Aldrich | Cat#: SML2351 |
| Acetaminophen | Sigma-Aldrich | Cat#: A7085 |
| Acetochlor | Sigma-Aldrich | Cat#: 33379 |
| Pymetrozine | Sigma-Aldrich | Cat#: 46119 |
| Clodinafop | Sigma-Aldrich | Cat#: 31676 |
| Fenoxycarb | Sigma-Aldrich | Cat#: 34343 |
| Thalidomide | Sigma-Aldrich | Cat#: T151 |
| Butyl benzyl phthalate | Sigma-Aldrich | Cat#: 36927 |
| Heptadecafluorooctane-sulfonic acid | Sigma-Aldrich | Cat#: 77282 |
| Methyl mercury | Sigma-Aldrich | Cat#: 442534 |
| Verapamil | Sigma-Aldrich | Cat#: V4629 |
| Carbamazepine | Sigma-Aldrich | Cat#: C4024 |
| Propiconazole | Sigma-Aldrich | Cat#: 45642 |
| Abamectin | Sigma-Aldrich | Cat#: 31732 |
| Benzylcetyldimethyl Ammonium Chloride | Sigma-Aldrich | Cat#: B4136 |
| Spirodiclofen | Sigma-Aldrich | Cat#: 33654 |
| Thiram | Sigma-Aldrich | Cat#: 45689 |
| Triadimefon | Sigma-Aldrich | Cat#: 45693 |
| Epoxiconazole | Sigma-Aldrich | Cat#: 36848 |
| Desmedipham | Sigma-Aldrich | Cat#: 45426 |
| Fenhexamid | Sigma-Aldrich | Cat#: 31713 |
| Cyproconazole | Sigma-Aldrich | Cat#: 46068 |
| Chlorpyrifos | Sigma-Aldrich | Cat#: 45395 |
| Propyzamide | Sigma-Aldrich | Cat#: 45645 |
| Thiacloprid | Sigma-Aldrich | Cat#: 37905 |
| Aspirin | Sigma-Aldrich | Cat#: A2093 |
| Permethrin | Sigma-Aldrich | Cat#: 45614 |
| Rotenone | Sigma-Aldrich | Cat#: 45656 |
| 5-methyltetrahydrofolic acid, disodium salt | Sigma-Aldrich | Cat#: M0132 |
| Dolutegravir | Sigma-Aldrich | Cat#: 10313 |
| Critical Commercial Assays |  |  |
| RPS18 Taqman Gene Expression Assay | Thermo-Fisher | Cat#Hs01375212_g1 |
| SuperScript IV First-Strand Synthesis System | Invitrogen | Cat#: 18091050 |
| Neon™ Transfection System | Invitrogen | Cat#: MPK10096 |
| Zymoclean Gel DNA Recovery Kit | Zymo Research | Cat#: D4007/D4008 |
| DC Protein Assay | Bio-rad | Cat#: 5000111 |
| Experimental Models: Cell Lines |  |  |
| WA09 hESC line (parent of all edited lines) | WiCell | (RRID:CVCL_9773) |
| Oligonucleotides |  |  |
| HoxA1 Taqman Gene Expression Assay | Thermo-Fisher | Cat#Hs00939046_m1 |
| HoxA5 Taqman Gene Expression Assay | Thermo-Fisher | Cat#Hs00430330_m1 |
| HoxC6 Taqman Gene Expression Assay | Thermo-Fisher | Cat#Hs00171690_m1 |
| HoxC8 Taqman Gene Expression Assay | Thermo-Fisher | Cat#Hs00224073_m1 |
| HoxC9 Taqman Gene Expression Assay | Thermo-Fisher | Cat#Hs00396786_m1 |
| HoxD10 Taqman Gene Expression Assay | Thermo-Fisher | Cat#Hs00157974_m1 |
| All SCRIB Taqman Gene Expression Assay | Thermo-Fisher | Cat#Hs00363005_m1 |
| SCRIB Iso A Taqman Gene Expression Assay | Thermo-Fisher | Cat#Hs01034954_g1 |
| SCRIB Iso B Taqman Gene Expression Assay | Thermo-Fisher | Cat#Hs01040361_g1 |
| Software and Algorithms |  |  |
| CellProfiler | Harvard | v4.2.6 |
| GraphPadPrism | GraphPad Prism | v10.2.1 |
| Nikon NIS-Elements (Acquisition) | Nikon | v5.42.01 |
| Nikon NIS-Elements AR (Analysis) | Nikon | v5.02.00 |
| RosetteDetect<sup>TM</sup> | Neurosetta LLC | v0.1.0.0 |
| Other |  |  |
| CryoMed controlled rate freezer | Thermo Fisher Scientific | Cat#: TSCM17PA |
| Prolong Gold Antifade Reagent | Invitrogen | Cat#: P36930 |
| Nikon A1R confocal microscope | Nikon | n/a |
| 4–20% Mini-PROTEAN® TGX<sup>TM</sup> Precast Gel | Bio-rad | Cat#: 4561094 |

### Lead Contact

Further information and requests for resources and reagents should be directed to and will be fulfilled by the Lead Contact, Randolph Ashton.

### Materials Availability

- Edited hPSC cell lines available upon request to lead contact.

### Data and Code Availability

- Any additional information required to reanalyze the data reported in this paper is available from the lead contact upon request.

#### Data

All data used for statistical analysis is included in the Supplemental Data excel file. All data reported in this paper will be shared by the lead contact upon request.

#### Code

- This paper does not report original code. RosetteDetect™ software is available at https://neurosetta.com/rosette-detect/ for use by the public.

### Experimental Model and Subject Details

#### Cell Lines

WA09 (H9, XX) hESC were supplied by the WiCell Research Institute. WA09 hESCs and the gene edited derivates were maintained at 37°C in 5% CO_2_ in E8 medium under feeder-free conditions on Matrigel (WiCell)-coated 6-well plates (Corning). For maintenance and expansion, the cells were subcultured in Versene every 5 days at a 1:12 ratio following ∼85% confluency.

For cell line authentication, each cell line (WA09 parent and 5 edited clones) was submitted to WiCell for karyotyping and mycoplasma testing (**Figure S14**).

### Method Details

#### RosetteArray Culture Substrate Fabrication

Culture substrates for 12-well RosetteArray assays were generated in accordance with our previously published protocol^21^. In brief, Polydimethylsiloxane (PDMS) stamps featuring rectangular arrays of circular recessions with 250, 150, 125, and 100 µm diameters were used to transfer and establish a self-assembled monolayer (SAM) of 2 mM ω-mercaptoundecyl bromoizobutyrate onto gold-coated No. 1 glass coverslips. Poly(ethylene) glycol methyl ether methacrylate (PEGMEMA) brushes were grafted from these SAMs using sodium ascorbate-initiated atom transfer radical polymerization (ATRP) under inert gas for 16 hours at room temperature. Trace copper ions and residual organic solvents were removed through subsequent washes with 70% ethanol. 96-well RosetteArray plates were obtained from Neurosetta LLC.

#### NMP Derivation

WA09 hESCs and gene-edited mutants were differentiated into NMPs as previously described^22,28^ utilizing CHIR99021, FGF8b, GDF11, and dorsomorphin. Briefly, hESCs were seeded at a density of 1.5 × 10^5^ cells/cm^2^ in E8 medium with 10 μM ROCK inhibitor for 24 hours. The medium was replaced with E6 medium on day 0 and then changed to E6 supplemented with FGF8b (200 ng/ml) 24 hours later (day 1). On day 2, cells were subcultured at a 2:3 ratio by washing once with PBS, incubating in Accutase for ∼2 minutes, and removing them from the surface with gentle pipetting. After centrifugation, cells were gently resuspended in NMP medium, i.e., E6 medium with FGF8b (200 ng/ml) and 3 μM CHIR99021, containing 10 μM Y27632 and seeded on Matrigel-coated plates. This initiates *HOX* colinear and combinatorial expression and is referred to as ‘Hox0’, with 0 representing the hours of CHIR exposure. NMP medium was replenished on day 4 (Hox48). On day 5 (Hox72), cervical spinal cells were collected for cryopreservation (see next section). Lumbar spinal cells used in the paper were taken directly from previously generated cryopreserved banks^22^ but they can be derived by caudalizing past the Hox72 time point. On day 5, cells should be subcultured (2:3) as before using NMP media. On day 7-10, the media should be replenished daily but now supplemented with GDF11 (30 ng/ml) and 1 μM dorsomorphin to stimulate caudal NMP development. On day 9, the cultures should be subcultured at a 1:1 ratio. On day 11 (Hox216), lumbar cells can be collected for cryopreservation.

#### Cryopreservation and Thaw of hESCs and NMPs

Confluent monolayers of WA09 hESCs in 6-well plates were prepared for cryopreservation through enzymatic dissociation in Accutase (ThermoFisher) and resuspend at ∼6,000,000 cells/mL in E8 medium with 10 µM ROCK inhibitor Y-27632 and 10% DMSO. WA09-derived NMPs were similarly dissociated in Accutase and resuspended in E6 medium at ∼6,000,000 cells/mL with 10 µM ROCK inhibitor and 10% DMSO. Cryopreservation was performed in Cryovials (Wheaton, Ref#: W985922) at 1mL/vial using a Thermo Scientific CryoMed Controlled-Rate Freezer (7450) set to lower temperature at the following rates; −10°C/min to 4°C, −1°C/min to −60°C, and −10°C/min to −100°C. Cryopreserved vials of cells were placed in liquid nitrogen dewars for extended storage. Cells were thawed at 37°C for 3-5 mins and resuspended in E8 medium with 10 µM ROCK inhibitor for hESC seeding or E6 medium with 10 µM ROCK inhibitor and 1 µM RA for NMP seeding.

#### Forebrain and Spinal Cord RosetteArray Derivation

FB RosetteArray derivation was initiated by thawing and seeding cryopreserved hESCs at ∼200,000 cells/cm^2^ onto Matrigel-coated micropatterned arrays with 250 µm diameter circular regions in E8 medium with 10 µM ROCK inhibitor Y-27632. After 1 day, they were cultured in E6 medium (with 0.1% DMSO when indicated) for 5-7 subsequent days using daily 50% media changes. SC RosetteArray derivation was initiated by thawing and seeding cryopreserved hNMPs^22^ at ∼150,000 cells/cm^2^ onto Matrigel-coated micropatterned arrays with 100 or 150 µm diameter circular regions in E6 medium, 10 µM ROCK inhibitor Y-27632, and 1 µM Retinoic Acid. After 1 day, they were cultured in E6 medium (with 0.1% DMSO when indicated) with 1 µM RA for 5 subsequent days using daily 50% media changes. Neuroectoderm monolayers were generated analogously in Matrigel-coated 6 well plates, without the use of micropatterned substrates. All culture was performed in tissue culture polystyrene plates (Corning) in medium supplemented with Penicillin/Streptomycin (ThermoFisher). For 96-well plate RosetteArray derivation experiments E6 medium supplementation with DMSO (Sigma-Aldrich) at 0.1% v/v was used.

#### RosetteArray DNT Testing

All DNT compound screening involved supplementation of the culture medium beginning 1 day after cell seeding and upon removal of ROCK inhibitor. Chemical compound stocks were prepared from dry stock, fresh for each experiment, and solubilized in DMSO or water. Dose response medium formulations were prepared through serial dilutions in E6 medium (with DMSO supplementation where indicated). Compound concentrations were doubled in formulations to account for 50% medium changes. Compounds solubilized in DMSO were normalized to control conditions containing DMSO supplementation matching the highest tested concentration of DMSO in the treatment groups. Enzymatic digestions of compounds for simulated metabolism experiments were performed with human S9 liver fraction. Compounds were digested at 1 mM concentrations with and without S9 liver fraction at 2 mg/mL for 1 hour at 37°C in 200 mM Tris buffer containing 40 mM NADPH. Digestions were terminated via aliquoting and freezing at −20°C. Dose response experiments with digested compounds were performed by daily thawing of fresh aliquots.

#### Immunocytochemistry

On the last day of culture, RosetteArrays were fixed in 4% paraformaldehyde (PFA) in PBS for ∼15 minutes. Micropatterned tissues were permeabilized and blocked with 0.1% Triton-X and 5% Donkey serum in PBS (PBS-DT) for 1 hour at room temperature. Primary antibodies listed in the Key Resource Table were incubated on the substrates at 4°C overnight and at a 1:200 dilution in PBS-DT. This was followed by 3 x 20-minute washes with PBS-DT. Next, secondary antibodies listed in Key Resources Table were incubated on the substrates at 4°C overnight and at a 1:500 dilution in PBS-DT. This was followed by 2 x 20-minute washes with PBS-DT, and cell nuclei were stained with DAPI for 10 minutes in the last PBS wash.

#### Image Acquisition

A Nikon AR-1 Scanning Confocal Microscope with an HD upgrade, Nikon DUG with GaAsP detectors, and a JOBS module software upgrade was used to acquire all fluorescent images. The JOBS module enables semi-automated image acquisitions. In each well, a 10x image was used to randomly selected tissues from the array. High definition, resonance or Galvano scanning image acquisition was performed using a 10, 20, or 60X objective (Nikon) at 512x512 (resonance) or 4096 x 4096 (Galvano) pixels to create Z-stacks with 5-7 image planes at 1.5 µm Z-axis spacing.

#### Image Analysis

Image segmentation (CellProfiler, Harvard) was used to quantify the number of DAPI^+^ and PAX6^+^ cells per image plane for each rosette. To minimize double counting of cells between image planes, only the middle image plane in a Z-stack was used to generate a relative number of cells (# of DAPI^+^ cells) and %PAX6^+^ (# of PAX6^+^ cells / # of DAPI^+^ cells) per tissue. Single rosette emergence was quantified through manual and RosetteDetect™ image analysis. For manual analysis, each tissue Z-stack was interrogated and scored as a binary single rosette structure (‘1’) or no or >1 rosette (‘0’). Neural rosettes in micropatterned tissues were identified by the presence of a coherent, polarized, N-cadherin^+^ or Phallodin^+^ ring structure. Percent neural rosette emergence was calculated as # single rosette-presenting vs. analyze tissues (n=40-50) per array. In single rosette tissues, rosette morphological characteristics were measured by outlining the boundary of the N-cadherin ring.

#### Calculation of Z’ Factor

The Z’ Factor was calculated as: Z’ = 1- (3(σ_postive_+ σ_negative_)/|μ_positive_ - μ_negative_|). Where σ_postive_ and σ_negative_ represent the positive (1.0 µM Methotrexate) and negative (E6/0.1% DMSO) controls’ standard deviation, and μ_positive_and μ_negative_are their respective means.

#### Calculation of the LOAEL

In the MTX dose-response experiment, %single neural rosette averages for each dose were compared to the experimental condition’s solvent control via a Student’s T-test within Excel. The first dose that was significantly different from the solvent control at a level of p<0.05 was considered the lowest-observed-adverse-effect level (LOAEL).

#### Non-linear Regression of Dose-response Curves

GraphPad Prism was used to calculate parameters associated with the Logistic Regression Model, plot sigmoidal dose-response curves, and analyze the curves’ fit for different parameters plus calculation of the 95% CIs and R^2^ values.

#### CRISPR/Cas9 gene editing

The UW-Madison Human Stem Cell Editing Core was used to create *MTHFR* and *SCRIB* gene-edited lines. sgRNA sequence identification for each editing site was completed using the CRISPOR design tool^86^. The sgRNAs were ordered from Synthego as a 1.5 nmol synthetic sgRNA with 2’-O-methyl 3’ phosphorothioate modification at the first and last 3 nucleotides following the recommended suggestion. hPSCs were cultured in TeSR-PLUS media (StemCell Technologies) on Matrigel until ∼80% confluency following standard cell culture protocols. Twenty-four hours before electroporation, cells were treated with CloneR (StemCell Technologies) following manufacturer protocol. Prior to the electroporation, the required sgRNA constructs were reconstituted following manufacturer protocols to a concentration of 150 pmole/µL. 1 µL of reconstituted sgRNA was pooled with 4 µg Cas9 Nuclease protein (TrueCut Cas9 Protein V2, Thermo Fisher Scientific) and 5 µL of Neon Buffer R (Invitrogen) to promote Cas9-RNP complex formation. After 15 minutes, 1.0 µL of ssODN primer (reconstituted to 1 µg/µL concentration, designed with homology overhangs of at least 40 base pairs) was added to the Cas9-RNP mix. Cells were singularized and lifted with a 1:1 mixture of 0.5 mM EDTA: Accutase for 3-4 minutes, resuspended in 1 mL PBS, and pelleted. Approximately 400,000 cells were resuspended in 35 µL Buffer R and mixed with 8 µL of the pre-prepared Cas9-RNP complex with repair ssODN. Cells were electroporated with a 10 µL NEON electroporation format using 1200V, 30 msec, 1x pulse settings. Cells were pooled following four rounds of electroporation and plated in TeSR-PLUS media with CloneR supplement at manufacturer-recommended concentrations following a serial dilution to promote single-cell clonal growth. Following expansion of 10-14 days, clones were identified and picked using standard techniques.

#### Genotyping

Bulk gDNA was collected from dissociated cells using QuickExtract DNA Extraction Solution 1.0 (Epicentre) to confirm editing efficiency prior to clonal selection. Single-cell clones were manually selected and mechanically disaggregated. Genomic DNA was isolated from a portion of these clones using QuickExtract DNA Extraction Solution 1.0. Genotyping primers were designed flanking the mutation site, allowing amplification of this region using Q5 polymerase-based PCR (NEB). PCR products were identified via agarose gel and purified using a Zymoclean Gel DNA Recovery Kit (Zymo Research). Clones were submitted to Quintara Biosciences for Sanger sequencing to identify clones with the proper genetic modification.

#### Off-target analysis

To identify whether the CRISPR-Cas9 system produced any non-specific genome editing, we analyzed suspected off-target sites for genome modification. Using the 5 highest-likelihood off-target sites for each sgRNA as predicted by the CRISPOR algorithms, we designed genotyping primers to amplify these regions via Q5-polymerase PCR. PCR products were identified via agarose gel, purified using a Zymoclean Gel DNA Recovery Kit, and submitted to Quintara Biosciences for Sanger sequencing.

#### qPCR and gene expression analysis

Total RNA was isolated from cell pellets using a TRIzol reagent (Invitrogen), and complementary DNA (cDNA) was synthesized using the SuperScript IV First-Strand Synthesis System (Invitrogen) according to the manufacturer’s instructions. TaqMan Gene Expression Assays and TaqMan Gene Expression Master Mix (Applied Biosystems) were used on a Bio-Rad CFX96 thermocycler with the following protocol: 50°C for 2 min; 95°C for 10 min; 40 cycles of 95°C for 15 s and 60°C for 1 min. Target genes were normalized to RPS18 expression, and relative gene expression was calculated using the comparative ▴Ct method. When fold differences (2^-▴Ct) are calculated and compared, RNA for each condition was collected in biologic triplicate and assayed in technical replicate for target genes. Values are reported for each gene with SDs after testing for the data’s lognormal distribution. Statistical analysis was conducted using GraphPad Prism software. Significance was determined using a one-way analysis of variance (ANOVA) with Tukey-Kramer post hoc test for multiple comparisons at a 95% confidence threshold.

#### Western Blot

SCRIB protein expression was qualitatively confirmed in the WA09 parent and P1043L mutant clones, and its absence was confirmed in the KO mutant clone via western blotting. Briefly, NEC monolayers derived in 6-well plates were pelleted and lysed in RIPA buffer (Thermo Fisher Scientific 89900) containing protease inhibitors (Thermo Fisher Scientific 78429) and stored at −80 °C. Upon thaw, lysates were analyzed for total protein concentration using the Lowry protein assay (Biorad DC Protein Assay). 25 µg of protein was analyzed on a 4-20% Mini-PROTEAN® TGX™ Precast Gel (Bio-Rad, 4561094) and transferred to polyvinylidene fluoride (PVDF) membranes (Thermo Scientific 88518). The membranes were cut to separate the 215 kDa SCRIB protein from the 50 kDA β-tubulin protein and blotted overnight at 4°C with an anti-SCRIB antibody (1:500; Invitrogen PA5-54821) or Anti-β-tubulin (BioLegend 1:500), respectively. Washes were conducted the next day, and membranes were incubated in horseradish peroxidase-conjugated goat anti-rabbit at room temperature, followed by addition of enhanced chemiluminescence (ECL) substrate (made in-house). Membranes were visualized using a chemiluminescence imager (DNR Bio-Imaging Systems) at 10 second intervals, ∼100s total exposure.

### Quantification and Statistical Analysis

GraphPad Prism (V10.2.1) was used for all statistical analysis. Error bars represent mean ± SD; * for p<0.05, ** for p<0.01, *** for p<0.001, **** for p<0.0001. Neural rosette emergence for single dose experiments were analyzed with t-tests and all other metrics were analyzed with One-way ANOVAs. Non-linear regression was performed to generate three (Hill Slope =1) or four (Hill Slope not restrained) parameter dose-response curves depending on whether data points were present at the curve’s inflection point. Comparison of dose-response curves for different metrics analyzing the same compound were compared using a hypothetical dose-response curve for all metrics. The explanation of “n” and the number of technical or biological replicates completed per experiment, as well as tests done to determine if data met assumptions of statistical approaches, can be found in the figure legends and text.

