## Supplemental Methods and Figures for "RosetteArray Platform for Quantitative High-Throughput Screening of Human Neurodevelopmental Risk"

**Supplemental File Inventory**

Within this file:

Additional uploaded files:

1. ‘Manuscript Data Tables’ excel file, which contains all data presented in both Figures and Supplemental Figures.
2. ‘Gene Editing Validation.zip’ file containing Sanger sequencing reads used to validate gene-edited clonal cell lines. (related to Figure S13)

### Supplemental Methods

**Derivation of RosetteArray-derived cortical organoids**

Cortical organoids were derived using custom 24-well RosetteArray^®^ plates (Neurosetta) which restrict cell adhesion to circular nodes with 250 µm diameter. On Day -2, RosetteArray plates were rinsed 3 times with DPBS (ThermoFisher) before the application of a solution of Matrigel in DMEM/F12 media (0.083 mg/mL protein concentration) with 0.5% Penicillin-Streptomycin (pen-strep, Gibco) and overnight incubation at 37^o^C.

On Day -1, cryopreserved hPSCs were thawed and resuspended in E8 media. To remove residual cryopreservation media, the cell suspension was then centrifuged at 200 x g before removing the supernatant and resuspending the cell pellet in fresh E8 media with 10µM ROCKi and 0.5% Pen-Strep. A sample of the cell suspension was then counted using trypan blue (ThermoFisher) on a Countess automated cell counter (ThermoFisher), and the cell suspension was diluted with E8+ROCKi to achieve a 125,000 cells/cm^2^ final seeding concentration. The Matrigel coating solution was then removed from the previously coated RosetteArray^®^ plates and 1mL cell suspension was dispensed into each well before placing the plate back into the 37^o^C incubator.

On day 0, plates were removed from the incubator and inspected for proper cell seeding. Then, 850µL of seeding media was removed from each well and replaced with 1mL of E6 media (DMEM/F12, 64 mg/L ascorbic acid, 543 mg/L sodium bicarbonate 14 ug/L sodium selenite, 19.4 mg/L insulin, 10.7 mg/L transferrin) + Pen-Strep. On Days 1 and 2, 50% media changes were performed by removing 500µL media from each well and replacing it with E6 + Pen-Strep.

Matrigel hydrogel overlays were applied on day 3. Plates were removed from the incubator, 850µL media was removed and 350µL Matrigel solution (at 50% concentration, diluted in E6 media) was added. Plates were then returned to the incubator for 30 minutes to facilitate gelation of the Matrigel hydrogel. 1mL of E6 was then added to each well and plates were returned to the incubator.

Daily 50% E6 media changes were performed on days 4-7. On day 8, 850µL E6 media was removed from each well and replaced with cortical differentiation media ((CDM), 50/50 base media of E6 and Neurobasal (ThermoFisher) with 0.5% (v/v) N2 supplement (ThermoFisher), 0.5% (v/v) MEM-NEAA (Gibco), 1% (v/v) B-27 supplement (ThermoFisher), and 1% (v/v) Glutamax (ThermoFisher)). Agitation was also initiated on day 8 by transferring plates to an orbital shaker at 100 rpm. Daily 50% media changes with CDM were performed on days 9-30.

On day 30, CDM was replaced with Neuroabasal+ media (Neurobasal media with 1x N2 supplement, 1x B-27 supplement, 1µM cAMP (Sigma), 10ng/mL GDNF (Peprotech), 10ng/mL NGF (Peprotech), 10ng/mL BDNF (Peprotech), 1x Glutamax, 0.5% (v/v) Pen-Strep). 50% media changes with Neurobasal+ were performed three times a week from day 30 to the end of the culture period.

**Whole-Mount Immunostaining and Tissue Clearing**

At the end of the desired culture period, cortical organoid tissues were rinsed once with DPBS before incubation at room temperature in 1mL/well of cold, 4% (w/v) paraformaldehyde solution (PFA). PFA was then removed, and each well was washed 3 x 10 minutes in DPBS. Tissues were then incubated in 1mL/well of 1.5% triton-x in DPBS (1.5% PBS-T) overnight at 4^o^C for permeabilization. The following day, tissues were incubated in a blocking solution of 5% donkey serum in 0.3% PBS-T (PBS-DT) for 2 hours. Primary antibodies were then diluted in PBS-DT, applied to cortical organoid tissues at 350µL/well, and incubated for 72 hours at 4^o^C. Primary antibodies were then removed and tissues were washed 5 x 30 minutes in 0.3% PBS-T before 24 hours of incubation in secondary antibodies (Alexa-fluor (ThermoFisher) diluted 1:500 in PBS-DT) with DAPI to label cell nuclei at 4^o^C. Tissues were then washed 5 x 15 minutes in 0.3% PBS-T. Tissue clearing was performed using a one-step clearing and mounting solution, RapiClear1.47 (SUNJIN Lab). Following washes, 350µL of RapiClear1.47 at 37^o^C was added to each well. Cortical organoids were then placed on a plate-rocker at a slow speed for 1 hour at room temperature, removed and placed at room temperature overnight, and then stored at 4^o^C before imaging.

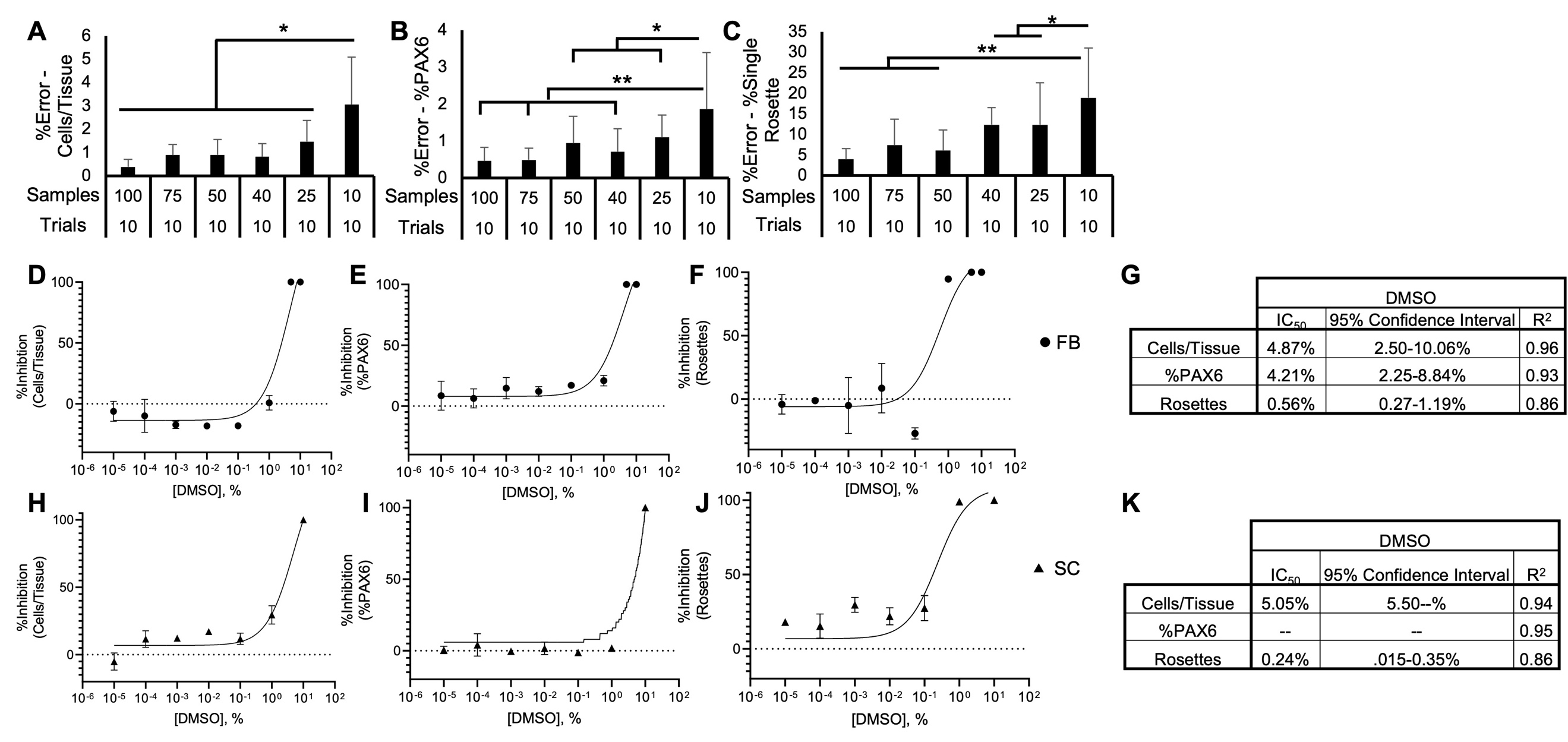

Figure S1. RosetteArray analysis sample size and %DMSO evaluations (related to Figure 1). Sampling different numbers of randomly selected tissues from a FB RosetteArray image dataset to determine %error when calculating (A) cell viability/proliferation, (B) neural induction, and (C) single rosette emergence. Error was calculated from 10 separate trials per sample number; stats conducted using a one-way ANOVA with Tukey-Kramer post-hoc analysis, *p ≤ 0.05, **p ≤ 0.01. DMSO dose-response (v/v%) for (D-G) FB (WA09 hPSC seeded) and (H-K) SC Rosette (cervical spinal NMP progeny seeded) arrays detailing %inhibition for (D, H) cells/tissue, (E, I) %PAX6, and (F, J) single rosette emergence with (G, K) descriptive stats for the non-linear regression, respectively. Each data point is the average of a WA09 hPSC or cervical spinal NMP progeny differentiation conducted in biological triplicate, n=50 technical replicates per well. Three-parameter non-linear regression used to model and compare each metric.

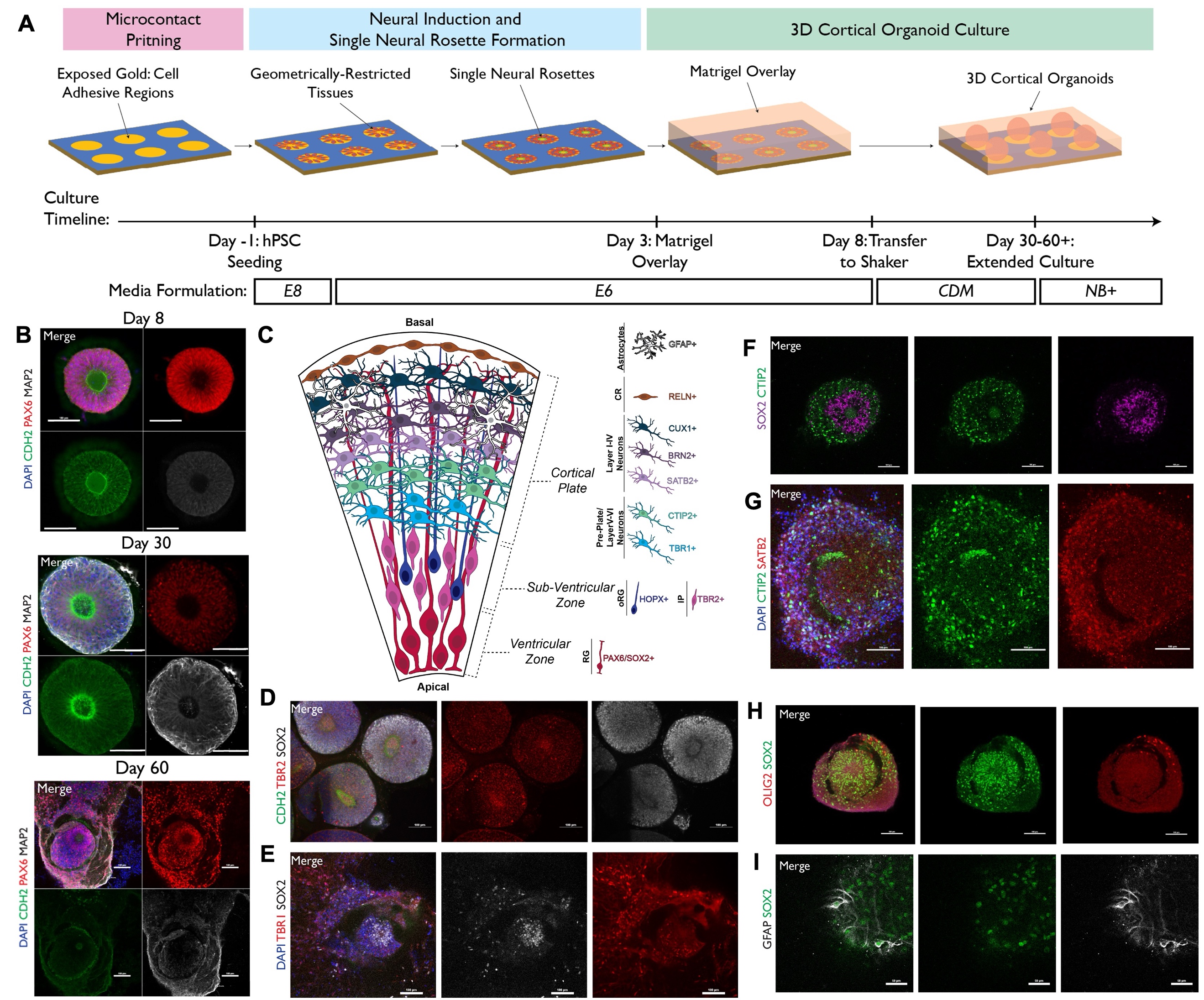

Figure S2. Cortical organoid derivation from micropatterned, singularly polarized, FB rosettes (related to Figure 1)**.** (A) Schematic of FB RosetteArray (WA09 hPSC seeded) long-term culture to generate arrayed cortical organoids. (B) Whole-mount immunostaining at Day 8, 30, and 60 organoids for Dapi (cell nuclei), N-cadherin (cell-cell adhesion), PAX 6 (neuroepithelial progenitors), and Map2 (post-mitotic neurons). (C) Schematic of cortical organoid radial organization with markers of progenitor and neuronal cell types. (D-G) Whole-mount immunostaining for cortical markers described in (C). Panel (D) is Day 16; (E,F) are Day 30; (**G**) are Day 60. Whole-mount immunostaining showing the presence of (H) oligodendrocyte precursors (Olig2+, Day 60) and (**I**) astrocytes (GFAP+, Day 47). Scale bars are (B,D-H) 100 µm and (I) 50 µm. Portions of (A,C) were created in Biorender.

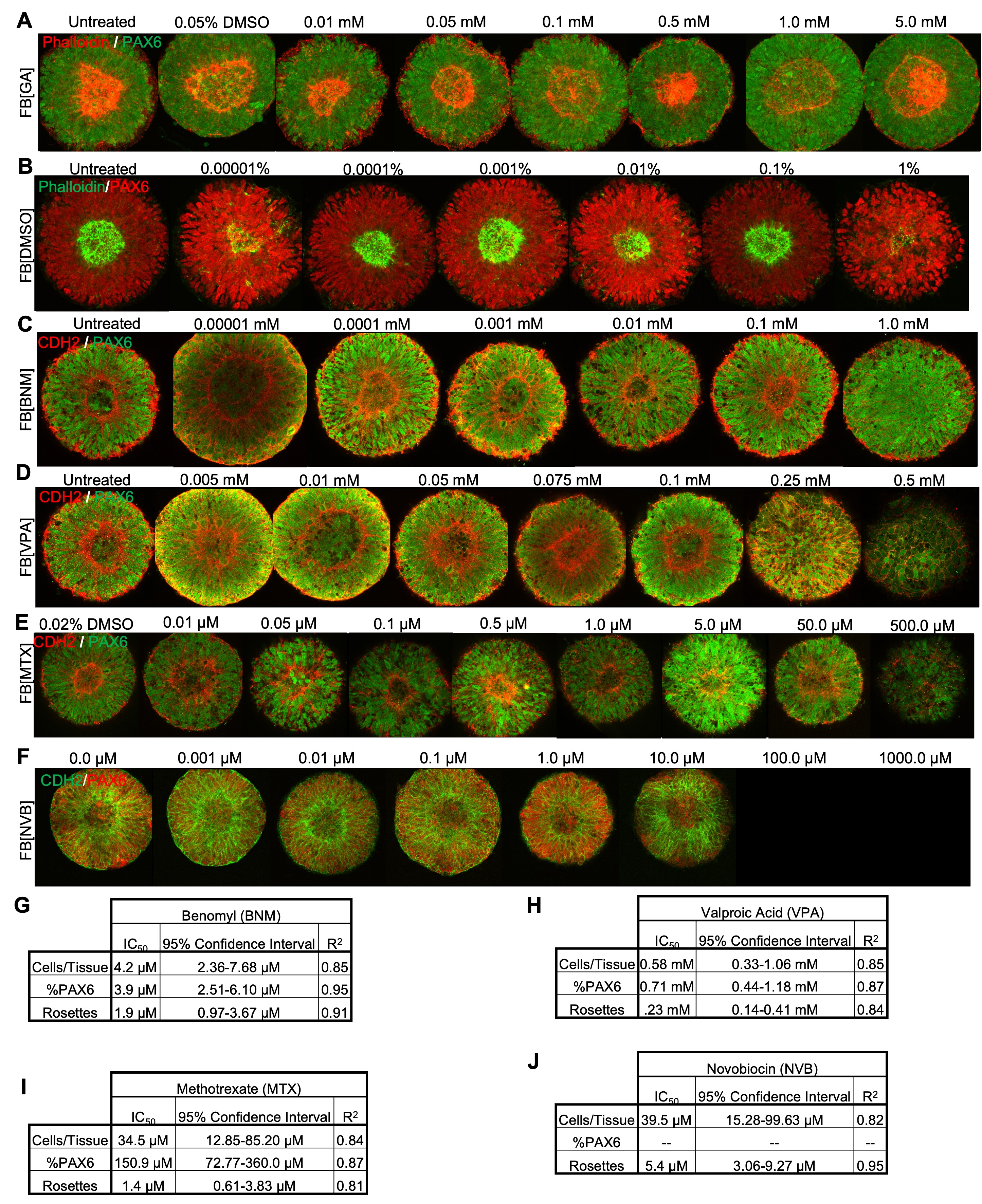

Figure S3. Corroborating data for FB RosetteArray dose-response experiments with NTD-risk chemicals (related to Figure 2)**.** Representative images of FB RosetteArray (WA09 hPSC seeded) tissues at each dose of (A) Glycolic acid, (B) DMSO, (C) Benomyl, (D) Valproic Acid, (E) Methotrexate, and (F) Novobiocin dose-response trials. (G-J), IC_50_ values and descriptive statistics for (G) Benomyl, (H) Valproic acid, (I) Methotrexate, and (J) Novobiocin dose-response experiments. In (A) and (B), Phalloidin (F-actin)^21^ is used to detect rosette polarization instead of N-Cadherin (CDH2). The DSMO concentration used across each dose response (i.e., equivalent to concentration at highest chemical exposure, except for DMSO dose-response) is indicated by the first image’s label with ‘Untreated’ or ‘0.0μM’ being E6 media with 0% DMSO. Scale bars are 100 µm.

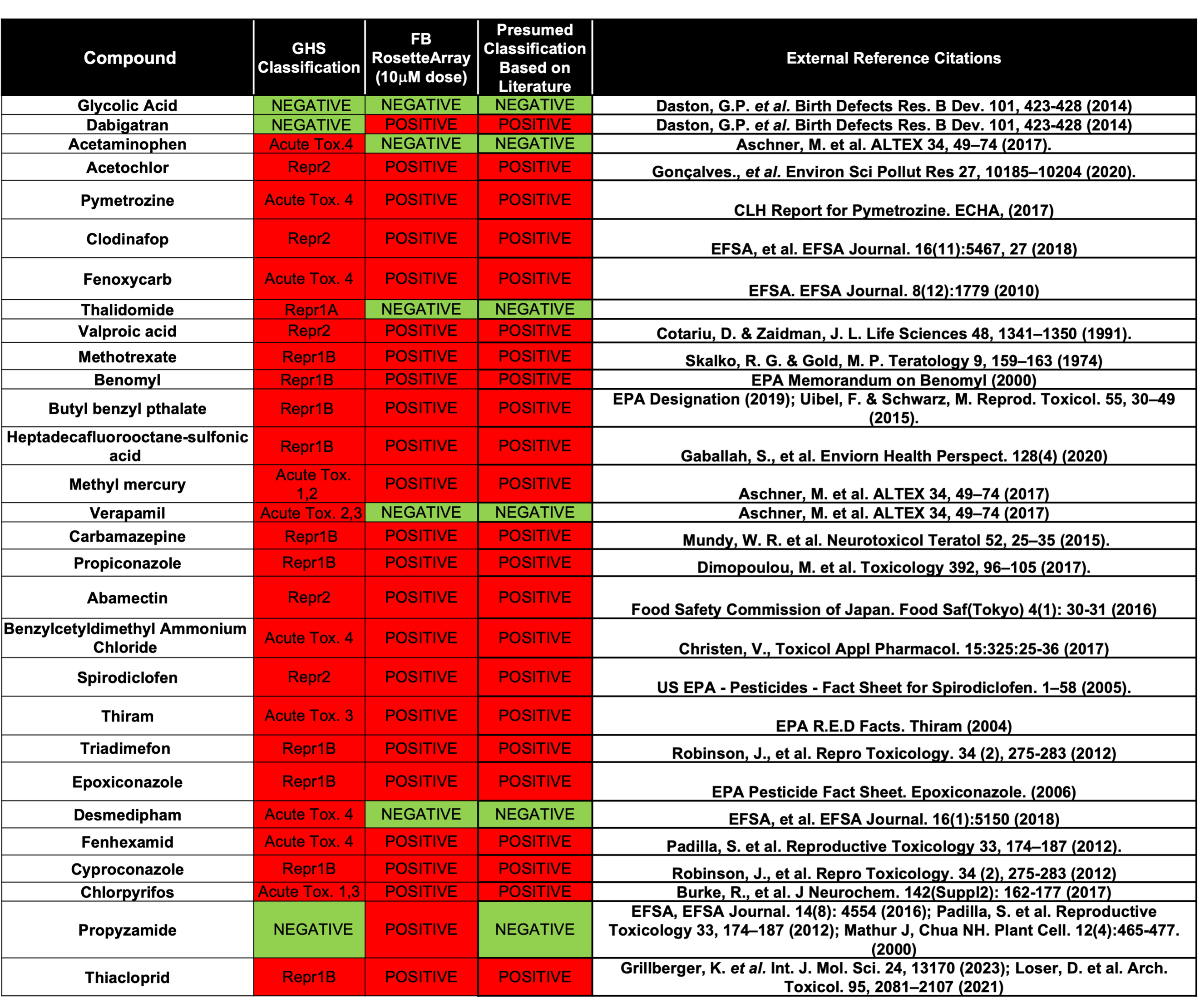

Figure S4. Compound classification references for DNT screen (related to Figure 4)**.**

Screened compounds with their Globally Harmonized System (GHS) and FB RosetteArray classification listed with literature reference citations for determination of ‘Positive’ or ‘Negative’ DNT classification.

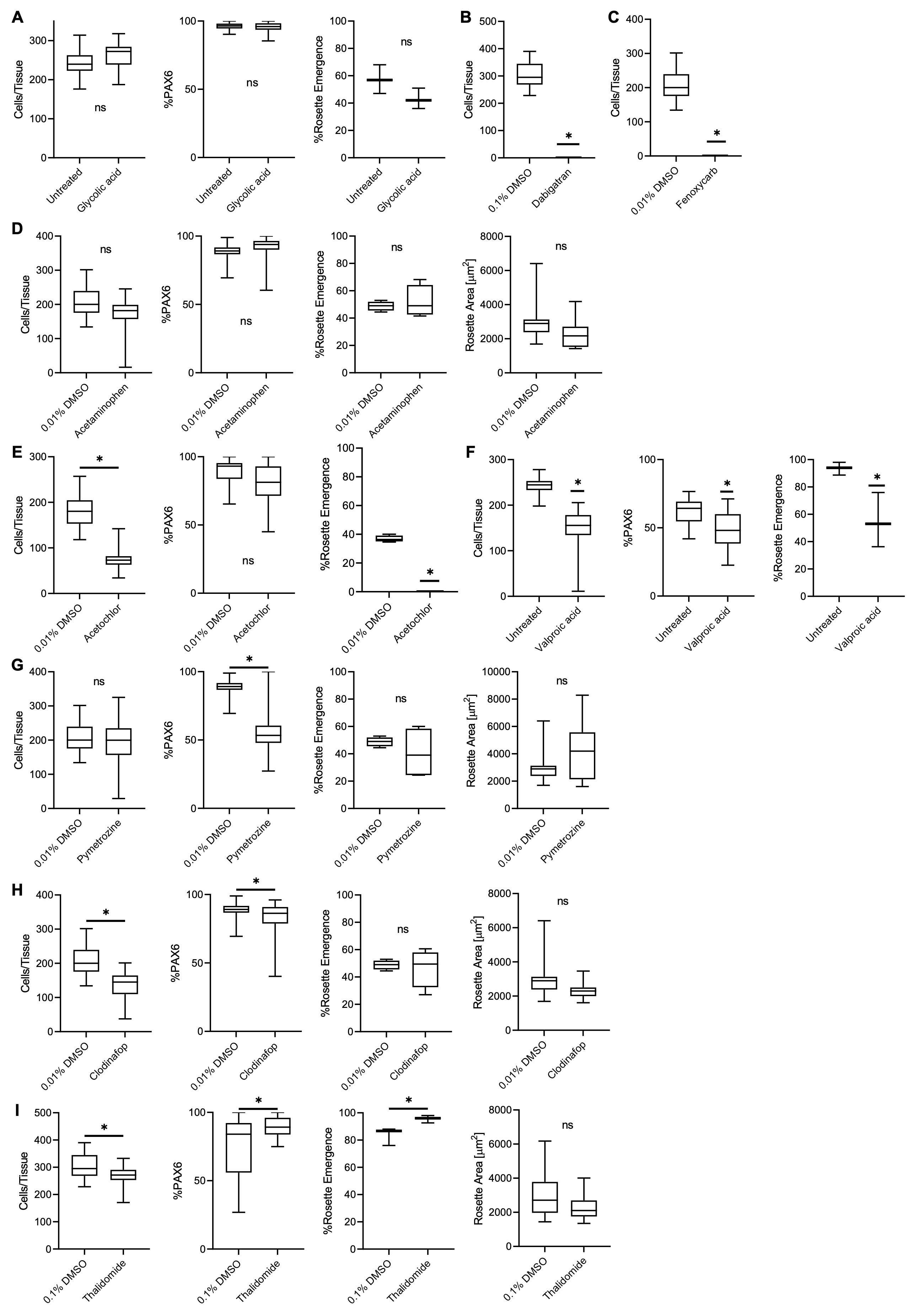

Figure S5. FB RosetteArray DNT screening plots (related to Figure 4)**.** Single dose (10 µM) toxicity experiments, compared to Untreated/media or % DMSO control with metrics represented in Figure 4, for (A) Glycolic acid, (B) Dabigatran, (C) Fenoxycarb, (D) Acetaminophen, (E) Acetochlor, (F) Valproic Acid, (G) Pymetrozine, (H) Clodinafop, and (I) Thalidomide. Each box plot represents a WA09 hPSC differentiation conducted in biological triplicate, n=50 technical replicates/well. Two-sample t test, *p ≤ 0.05.

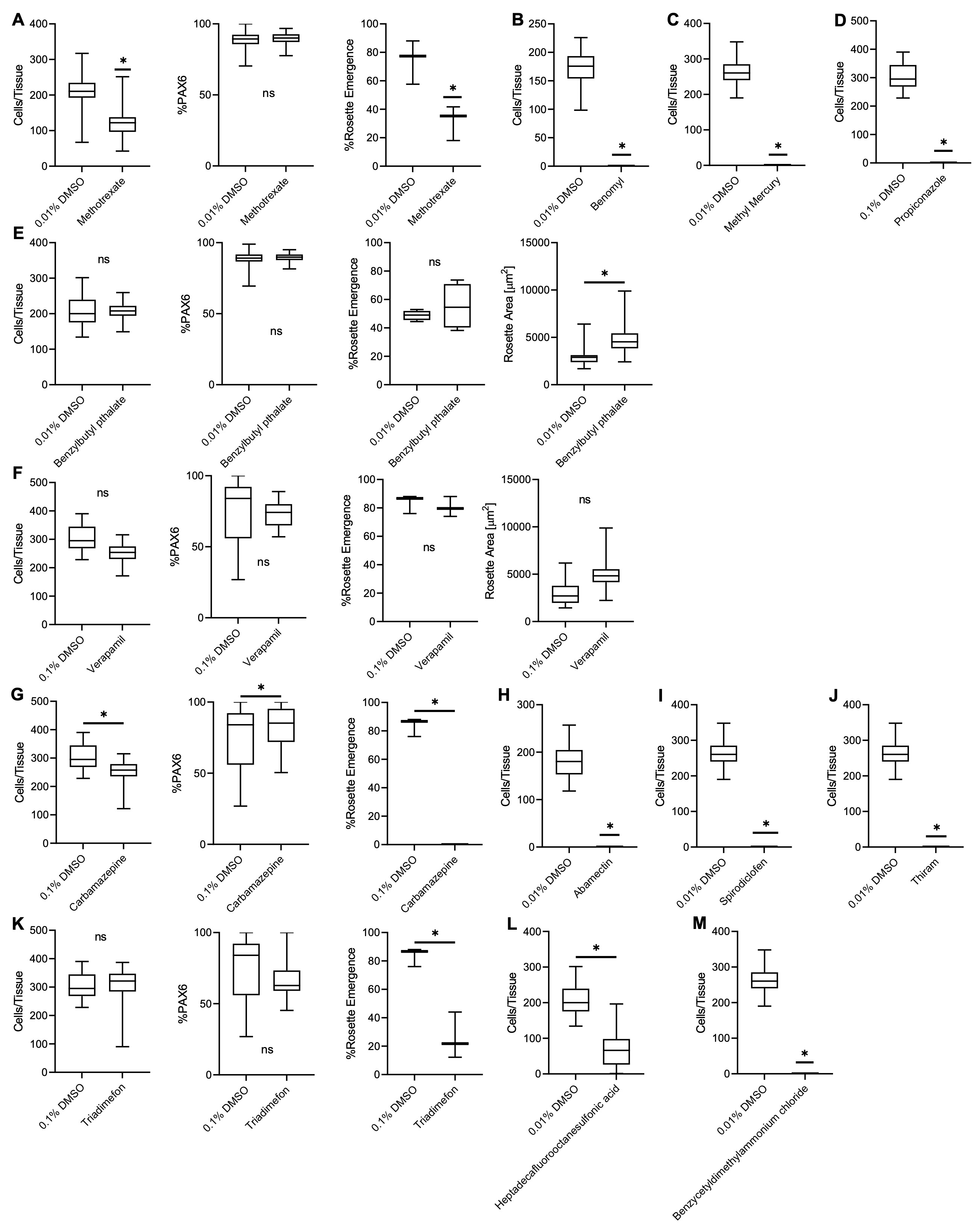

Figure S6. FB RosetteArray DNT screening plots (cont.) (related to Figure 4)**.** Single dose (10 µM) toxicity experiments, compared to % DMSO control with metrics represented in Figure 4, for (A) Methotrexate, (B) Benomyl, (C) Methyl Mercury, (D) Propiconazole (E) Benzyl butyl phthalate, (F) Valproic Acid, (G) Carbamazepine, (H) Abamectin, (I) Spirodiclofen, (J) Thiram, (K) Triadimefon, (L) Heptadecaflurooctanesulfonic acid, and (M) Benzycetyldimethylammonium chloride. Each box plot represents a WA09 hPSC differentiation conducted in biological triplicate, n=50 technical replicates/well. Two-sample t-test, *p ≤ 0.05.

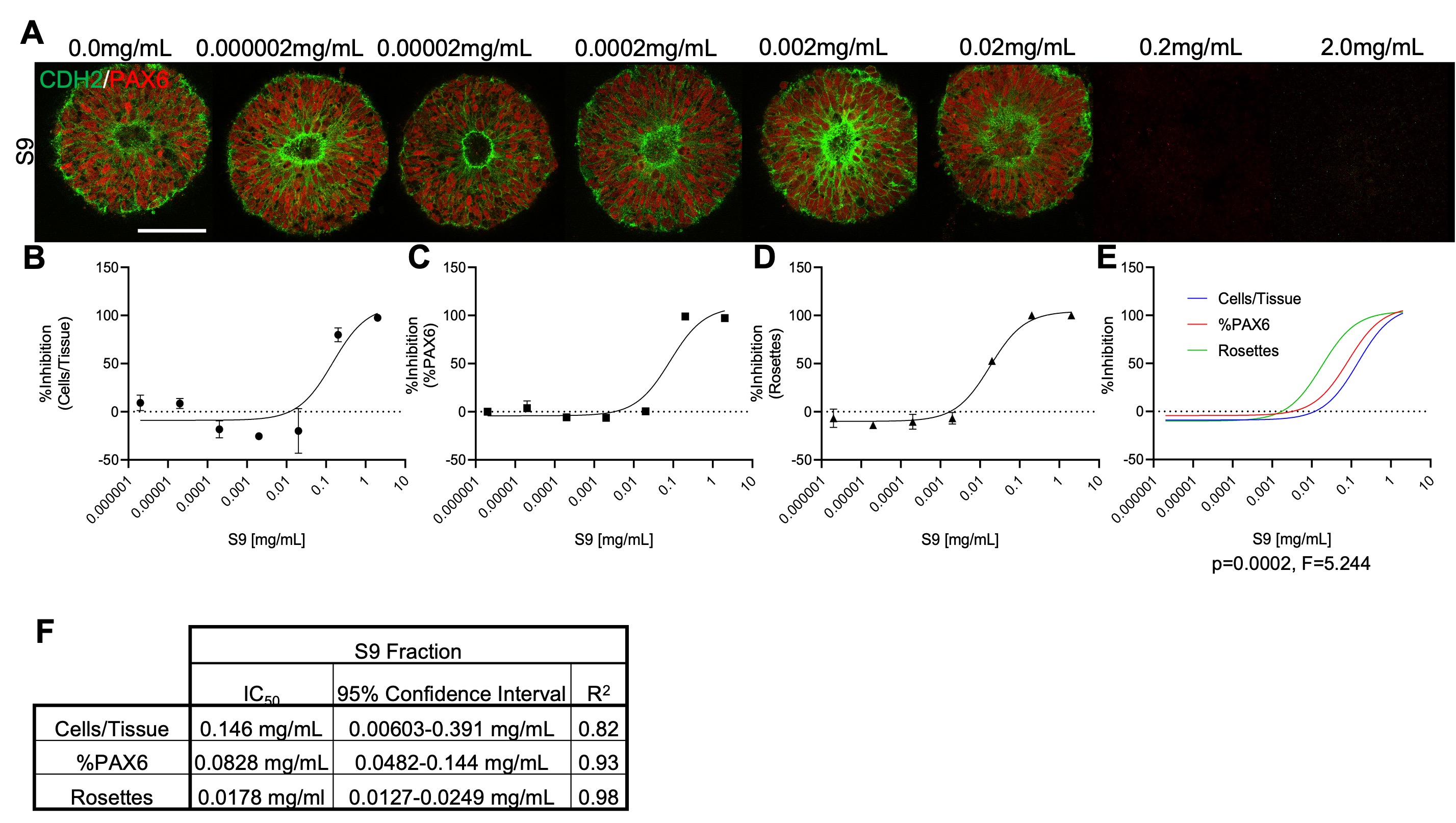

Figure S7. FB RosetteArray S9 dose-response images and plots (related to Figure 4)**.** (A) Representative immunostaining of dose-response experiments with human S9 liver fraction quantified for (B) cell viability/proliferation, (C) %PAX6, and (D) single rosette emergence. Each data point is the average of a WA09 hPSC differentiation conducted in biological triplicate, n=50 technical replicates/well. (F) Descriptive statistics for S9 dose-response curves. 0.1% DMSO media supplementation was used in these experiments. Scale bar is 100 µm.

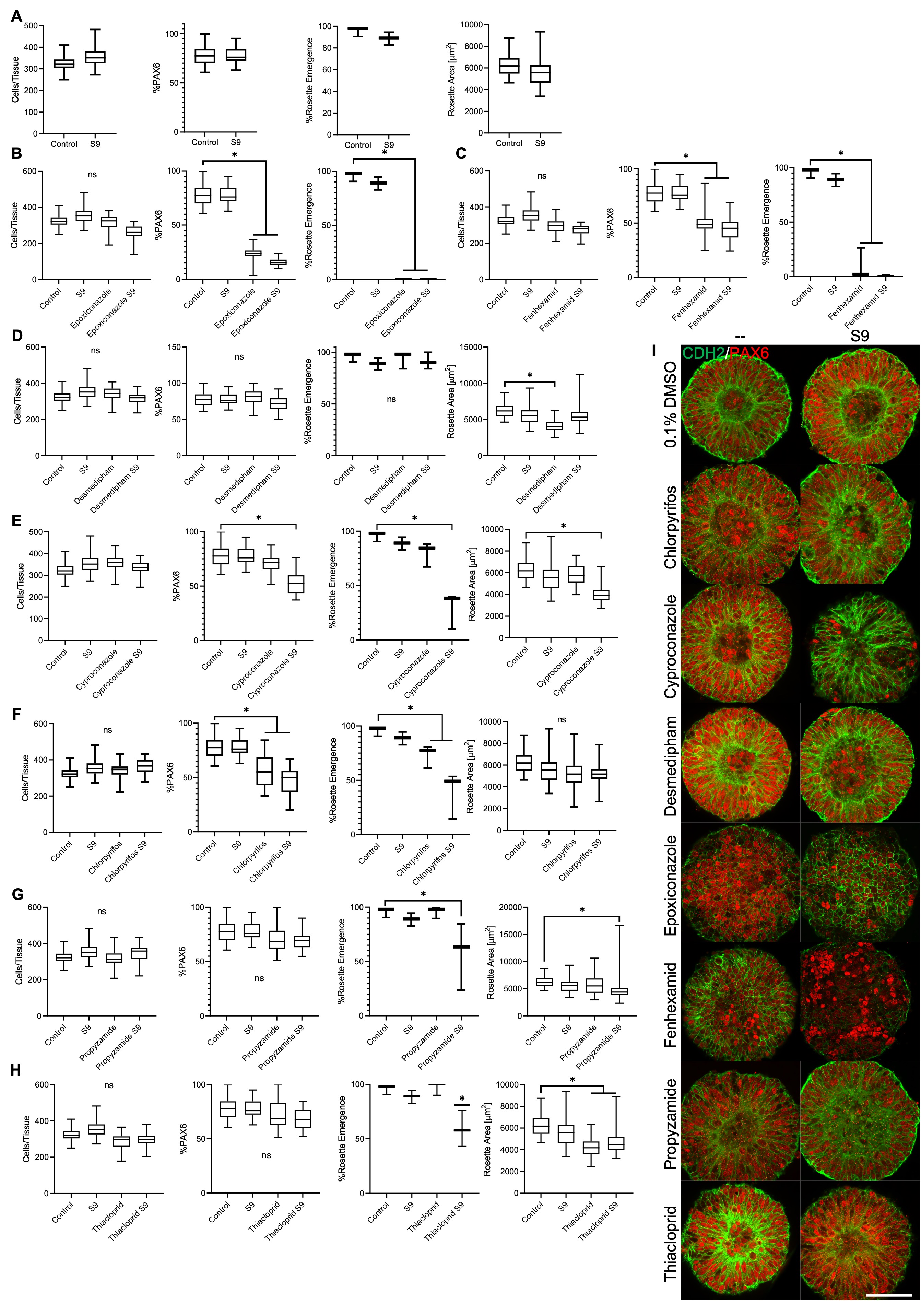

Figure S8. FB RosetteArray DNT screening with simulated human metabolism plots (related to Figure 4)**.** Single dose (10 µM) toxicity experiments with and without S9 metabolism, compared to 0.1% DMSO media supplementation (‘Control’) with metrics represented in Figure 4, for (A) 0.1% DMSO control, (B) Epoxiconazole, (C) Fenhexamid, (D) Desmedipham (E) Cyproconazole, (F) Chlorpyrifos, (G) Propyzamide, and (H) Thiacloprid. (I) Representative fluorescent images of FB RosetteArray tissues exposed to various compounds with and without S9 pre-digestion. Each box plot represents a WA09 hPSC differentiation conducted in biological triplicate, n=50 technical replicates/well. One-way ANOVAs, *p ≤ 0.05.  Scale bar is 100 µm.

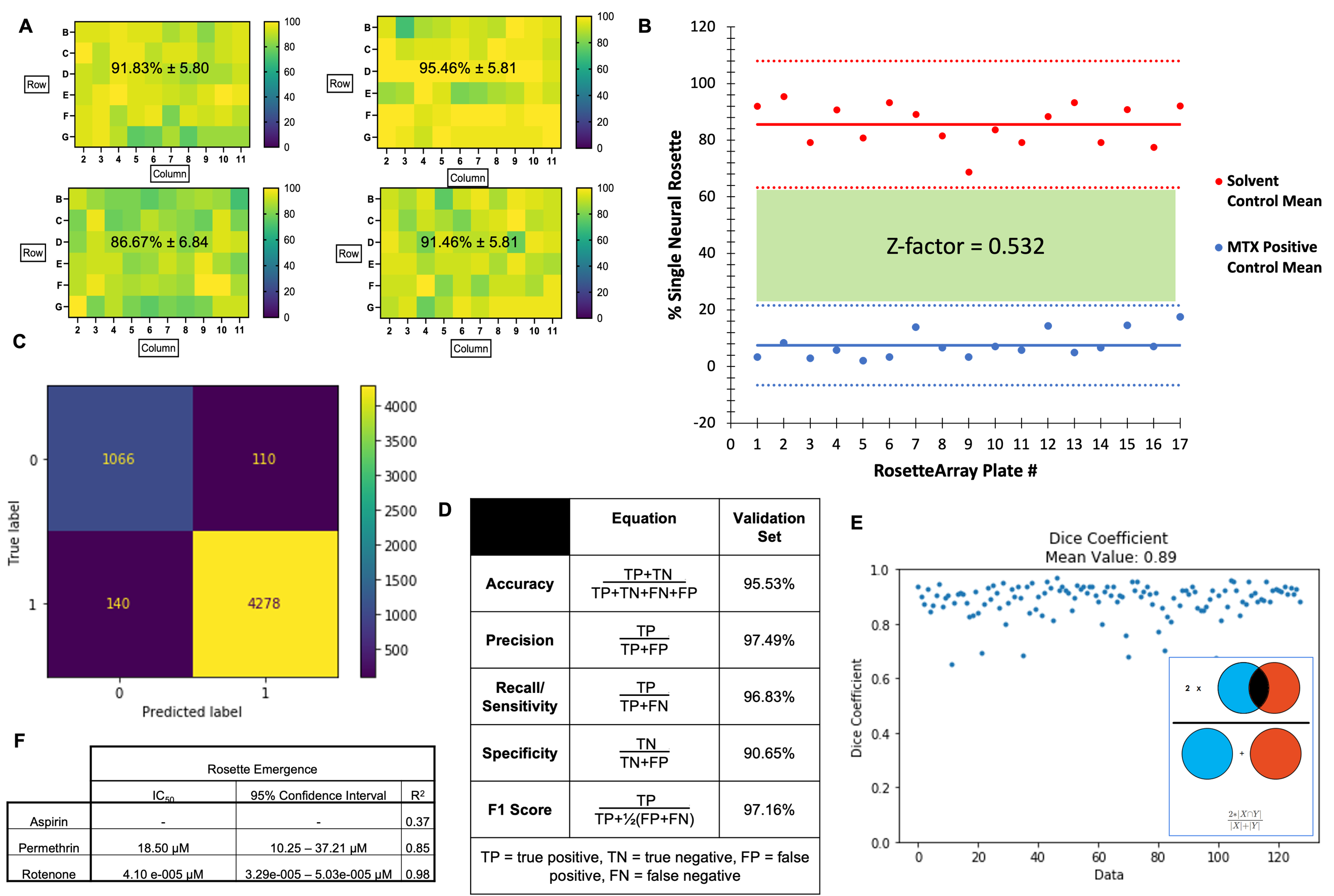

Figure S9. 96-well FB RosetteArray scale-up and RosetteDetect validation (related to Figure 5)**.** (A) Heat map of single neural rosette emergence efficiency across the interior 60 wells for four replicates of 96-well FB RosetteArrays using direct seeding of cryopreserved WA09 hESCs. (B) Z-factor analysis of 17 FB RosetteArray plates comparing single rosette emergence values/averages for E6/0.1% DMSO solvent controls and 1.0 μM MTX positive controls. Each data point represents the average of a WA09 hPSC differentiation conducted in 3-6 biological replicates, n=40 technical replicates per well. (C) Confusion matrix determined from RosetteDetect test image dataset post training. (D) Table of RosetteDetect performance metrics across the test image dataset. (E) Dice coefficient graph showing concordance between manually vs. RosetteDetect curated segmentation of polarized N-cadherin^+^ area across 128 test images. (F) Table of calculated IC_50_ doses, 95% confidence interval metrics, and R^2^ for Aspirin, Permethrin, and Rotenone FB RosetteArray dose-response curves.

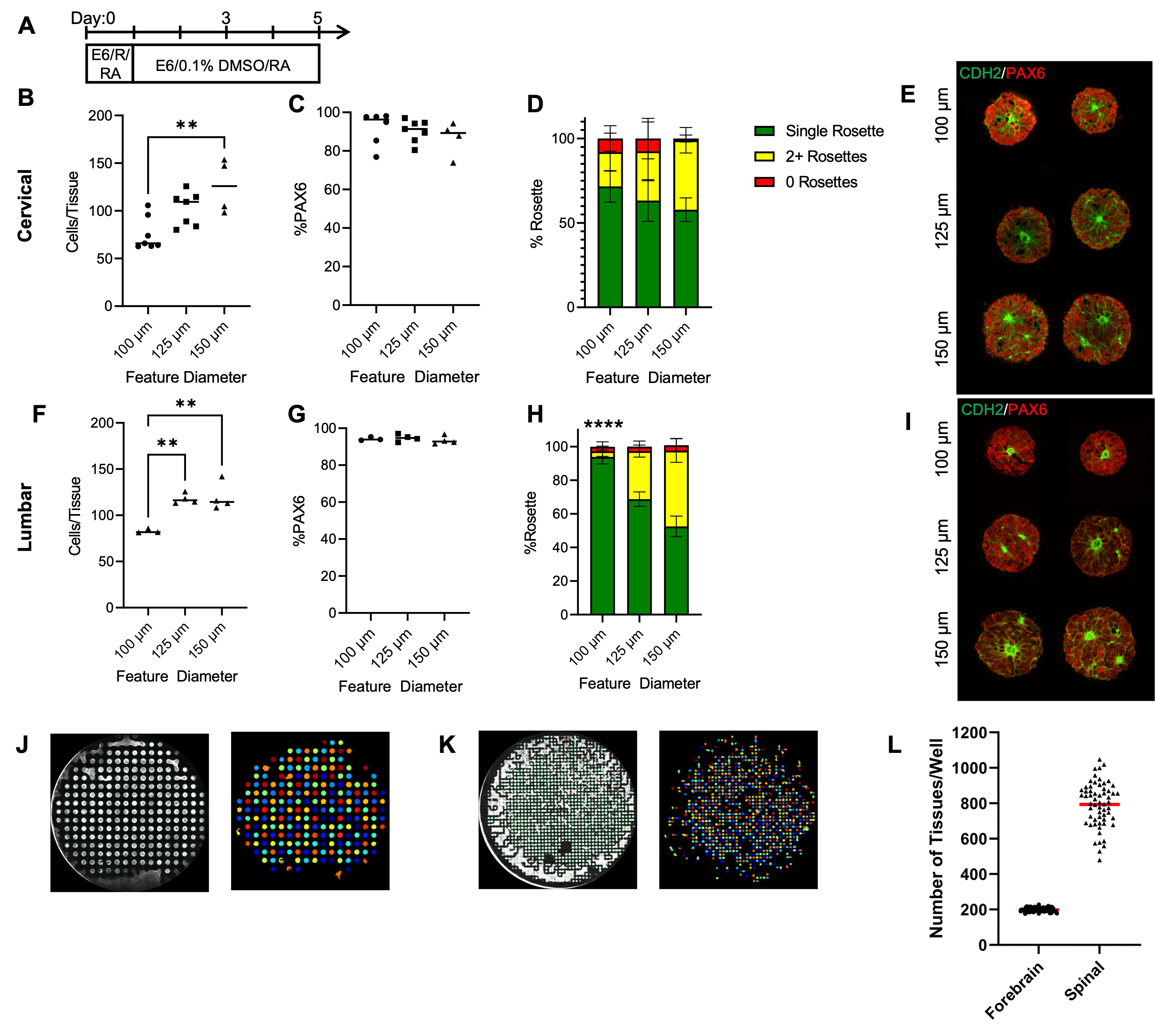

Figure S10. SC RosetteArray optimization (related to Figure 5) and tissue per well quantification**.** (A) Culture schema of 12-well plate RosetteArray format used for direct seeding of cryopreserved cervical and lumbar spinal NMPs onto micropatterned substrates. Quantification of cells/tissue, %PAX6, and %single rosette emergence with representative images for varied feature diameters for (B-E) cervical and (F-I) lumbar SC RosetteArrays, respectively. Each data point and bar graph represents a cervical or lumbar NMP differentiation conducted in 3-7 biological replicates with n=50 technical replicates per well. (D, H) Bar graphs of rosette emergence quantification showing distribution of tissues with 0, 1, and 2+ rosette structures. Representative images of micropatterned tissues within (J) FB and (K) lumbar SC RosetteArray 96-well plates with (L) tissue per well quantification across a single plate (n=60 wells; FB mean= 198.9 ± 11.6; SC mean= 793.6 ±125.7) using CellProfiler™. SC RosetteArray plate was slightly over-seeded causing loss of physical tissue separation around the well border. Statistical significance determined using a one-way ANOVA with Tukey-Kramer post-hoc analysis **p ≤ 0.01, ****p ≤ 0.0001.

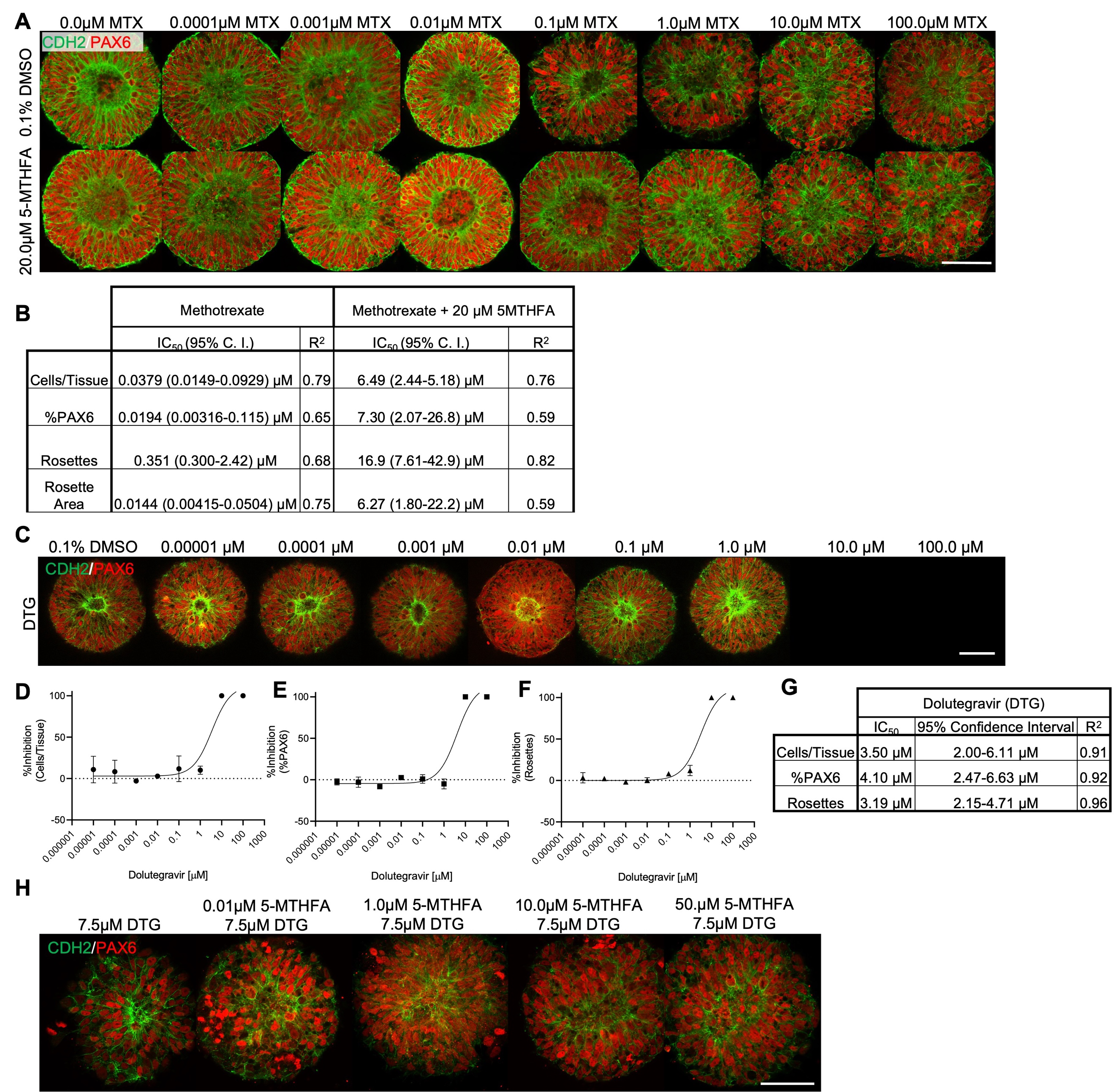

Figure S11. Folate metabolic pathway-specificity experimental images, dose-response curves, and descriptive stats (related to Figure 6)**.**  (A) Confocal images of representative 12-well FB RosetteArray tissues at each MTX concentration with and without 5-MTHFA supplementation. (B) Descriptive stats for dose-response curves displayed in Figure 5C-F. (C) Representative images of DTG dose-response on 12-well FB RosetteArraywith %inhibition curves of (E) cells/tissue, (E) neural induction, and (F) single rosette formation with (G) descriptive statistics for each curve. Each data point represents a WA09 hPSC differentiation conducted in biological triplicate, n=50 technical replicates per well. (H) Representative images of 7.5 μM DTG-exposed FB RosetteArray tissues with increasing 5-MTHFA concentrations. 0.1% DMSO media supplementation was used in these experiments. Three-parameter non-linear regression used to model and compare each metric. Scale bars are 100 µm.

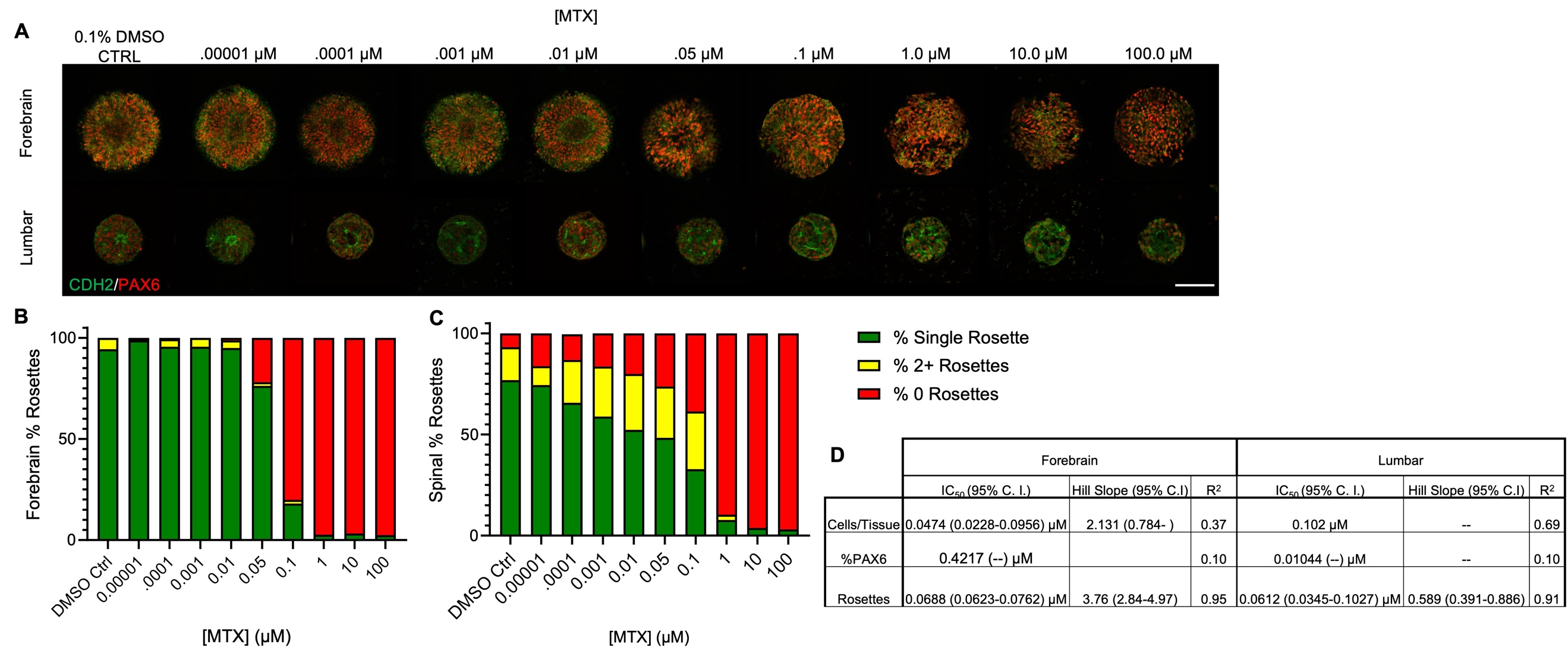

Figure S12. 96-well FB and SC RosetteArray-MTX dose-response images and descriptive stats (related to Figure 6)**.** (A) Confocal images of representative 96-well FB and lumbar SC RosetteArray tissues at each concentration of MTX. %Rosette counts per tissue at each dose of MTX for (B) FB and (C) lumbar SC Rosette Arrays. Each bar graph represents a WA09 hPSC or lumbar spinal NMP progeny differentiation conducted in biological quadruplicate, n=40 technical replicates per well. (D) Descriptive stats for curves displayed in Figure 6M-O. 0.1% DMSO media supplementation was used in these experiments. Scale bars are 100 µm.

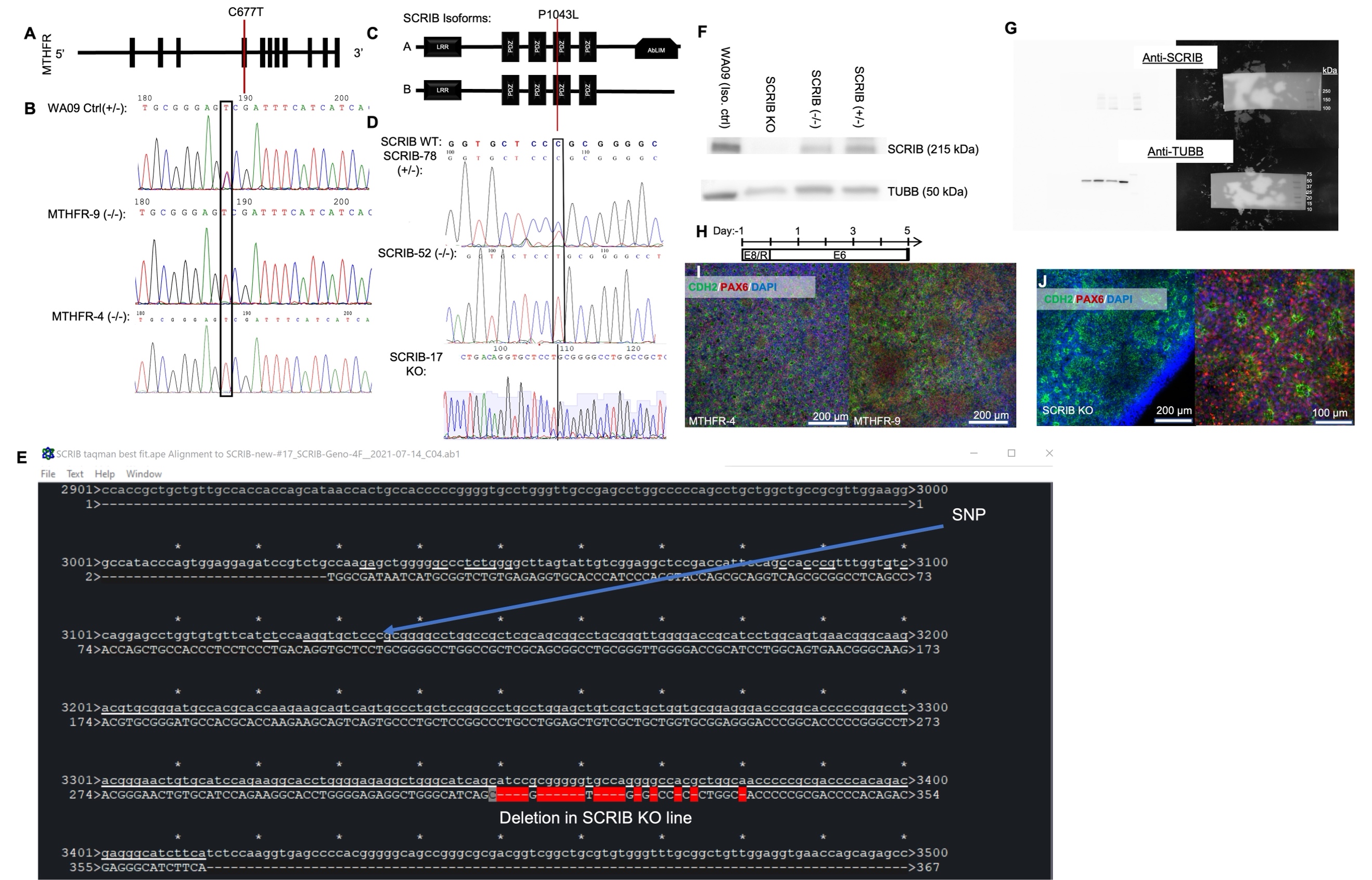

Figure S13. Analysis of MTHFR and SCRIB gene-edited lines (related to Figure 7)**.** (A) Representation of coding regions in human MTHFR gene with C677T SNP labeled. (B) Sanger sequence of homozygous MTHFR-#4 and -#9 mutant clones and the heterozygous WA09 isogenic parent line. (C) Representation of human SCRIB protein isoforms with P1043L mutation labeled. (D) Sanger sequence of the (+/-), (-/-), and KO gene-edited clones for c.C3128T SNP. (E) Sequence representation of nonsense deletion in the SCRIB KO line with (F) qualitative protein expression from rosette-forming NECs from each mutant line and the WA09 parent line. (G) Raw western blot images; reverse lane order from (F). (H) Culture schema for 6-well plate E6 neural differentiation. (I, J) Representative immunostaining of NECs derived from MTHFR-4 and -9 and the SCRIB^KO^ lines.

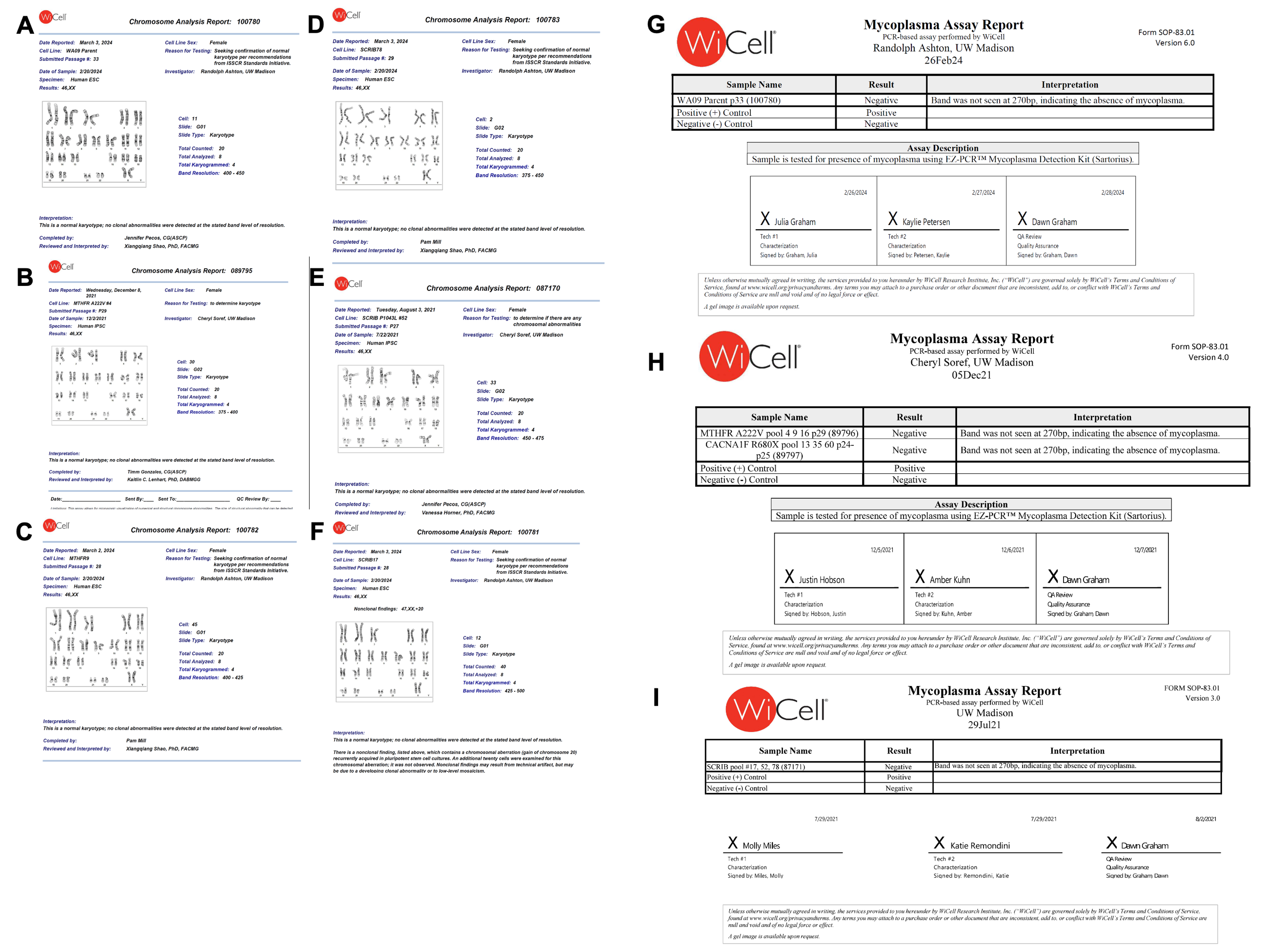

Figure S14. Cell line karyotypes and mycoplasma testing (related to Figure7)**.**  Representative karyotypes of (A) WA09 parent, (B) MTHFR-4^(-/-)^, (C) MTHFR-9^(-/-)^, (D) SCRIB-78^(+/-)^, (E) SCRIB-52^(-/-)^, and (F) SCRIB-17^KO^ lines. Pooled mycoplasma testing for (G) WA09 parent control, (H) MTHFR, and (I) SCRIB lines. All conducted by WiCell.

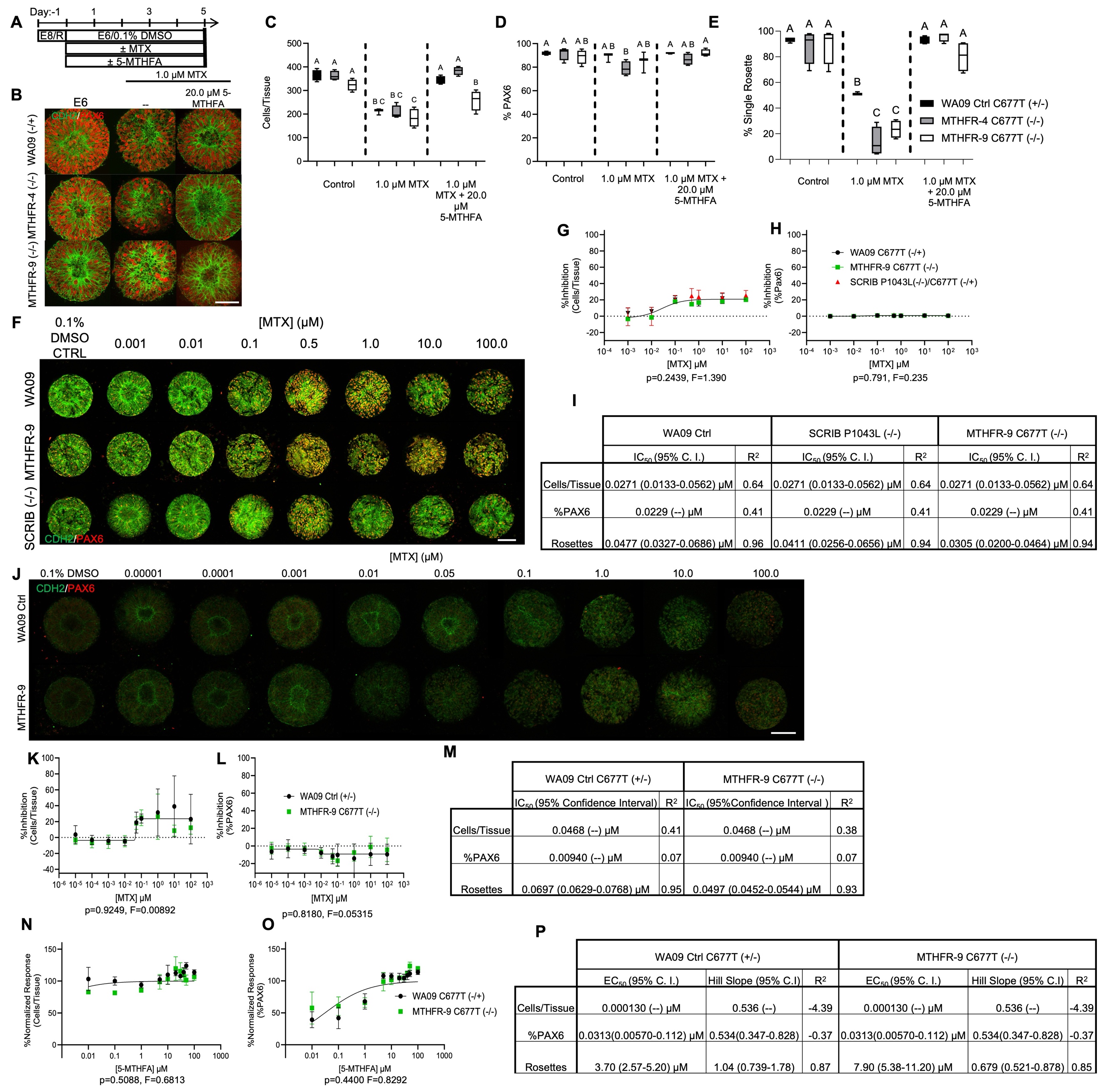

Figure S15. FB RosetteArray detection of folate metabolic pathway-specific genetic NTD risk: preliminary multifactorial scenario data, experimental images, dose-response curves, and descriptive stats (related to Figure 7)**.**  (A) Culture schema for FB RosetteArray 20 mM 5-MTHFA rescue of MTHFR^C677T (-/-)^ mutant lines and their MTHFR^C677T (+/-)^ WA09 parent control under 1 μM MTX with (B) representative staining and quantification of (C) Cells/Tissue, (D) %PAX6, (E) %single rosette emergence. Each box plot represents a WA09 or mutant line differentiation conducted in biological quadruplicate, n=50 technical replicates per well. Significance assessed using one-way ANOVA with Tukey-Kramer post-hoc analysis at p ≤ 0.05. (F) Representative immunostaining, %inhibition curves of (G) Cells/Tissue and (H) %PAX6, and (I) descriptive stats for these curves and the %single rosette emergence curve from Fig. 7B. (J) Representative immunostaining, %inhibition curves of (K) Cells/Tissue and (L) %PAX6, and (M) descriptive stats for these curves and the %single rosette emergence curve from Fig. 7C. %Normalized response curves of (N) Cells/Tissue and (O) %PAX6, and (P) descriptive stats for these curves and the % normalized single rosette emergence curve from Fig. 7F. Each data point on a dose-response curve represents a WA09 or mutant hPSC lines differentiation conducted in biological quadruplicate, n=40 technical replicates per well. Four-parameter non-linear regression used to model and compare each metric. If curves are statistically equivalent, then only one curve is displayed. Scale bars are 100 µm.

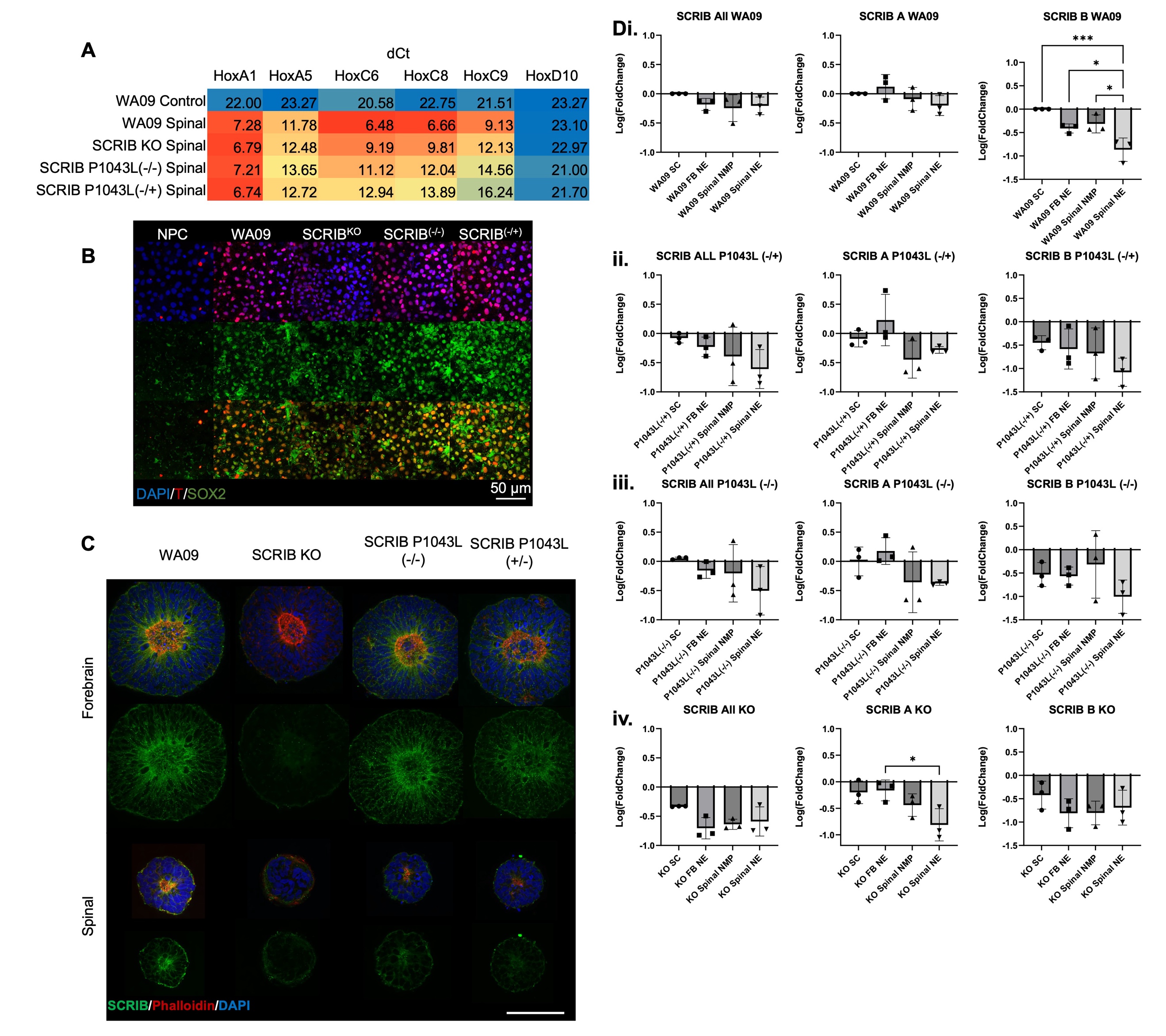

Figure S16. FB and cervical SC RosetteArray detection of PCP pathway genetic NTD risks: qPCR and immunostaining data (related to Figure 7)**.**  (A) Delta-Ct values for HOX gene expression and (B) Brachyury (T)/Sox2 co-staining, with no primary control (NPC), in cervical spinal NMPs used to generate RosetteArrays in Figure 7G-N. (C) Additional representative immunostaining for SCRIB and F-actin (Phalloidin) in FB and SC rosette tissues. (D) qPCR of stem cell (SC), FB neuroectoderm (NE), spinal NMP, and spinal neuroectoderm (NE) cell states for total (all) SCRIB and isoform A/B expression across the (i) WA09 parent, (ii) SCRIB^P1043L(+/-)^, (iii) SCRIB^P1043L(-/-)^, and (iv) SCRIB^KO^ lines normalized to WA09 SC expression levels. Each bar graph data comes differentiating a WA09 or mutant hPSC line in biological triplicate with each data point being the average of qPCR conducted in technical duplicate, and distribution confirmed to be log-normal via GraphPad Prism analysis. Comparison via one-way ANOVA with Tukey-Kramer post-hoc analysis, *p ≤ 0.05 and ***p ≤ 0.001. Scale bar is 100 µm.
